# Diversity of pheromone temporal coding disruptions by plant volatiles

**DOI:** 10.64898/2026.03.24.713794

**Authors:** Paul Clémençon, Tomáš Bárta, Christelle Monsempès, Michel Renou, Philippe Lucas

## Abstract

Moth pheromone-sensitive olfactory receptor neurons (Phe-ORNs) encode the intermittent structure of pheromone plumes through precisely timed spikes, a mechanism that is essential for odor plume tracking behavior in flying insects. However, natural olfactory scenes are composed of diverse volatile plant compounds (VPCs) with complex temporal dynamics whose effects on pheromone signal intermittency encoding remain unclear. Two lines of research, encoding of pheromone intermittency and background interference, remain largely disconnected. Here, we performed electrophysiological recordings from moth Phe-ORNs to quantify their responses to turbulent plume-like flickering pheromone stimuli in constant or fluctuating backgrounds of a diversity of VPCs. We found that some VPCs reversibly disrupted the temporal coding of various subregions of the pheromone stimulus and the trial-to-trial variability. While Phe-ORN activation by VPCs partially accounted for the decrease in coding performance, Phe-ORN gain reduction was insufficient to explain the full extent of the disruption. Some VPCs disrupt temporal coding without activating Phe-ORNs, and others activate Phe-ORNs without altering temporal coding. A continuous background noise can induce strong adaptation and limit dynamic range, whereas a fluctuating background can interfere with pheromone pulse encoding by disrupting spike timing. Altogether, these results indicate that the pheromone detection system must contend with multiple forms of background noise rather than a uniform disturbance. Timing is key for olfactory navigation, and our results raise questions regarding how downstream circuits would process noisy sensory inputs.

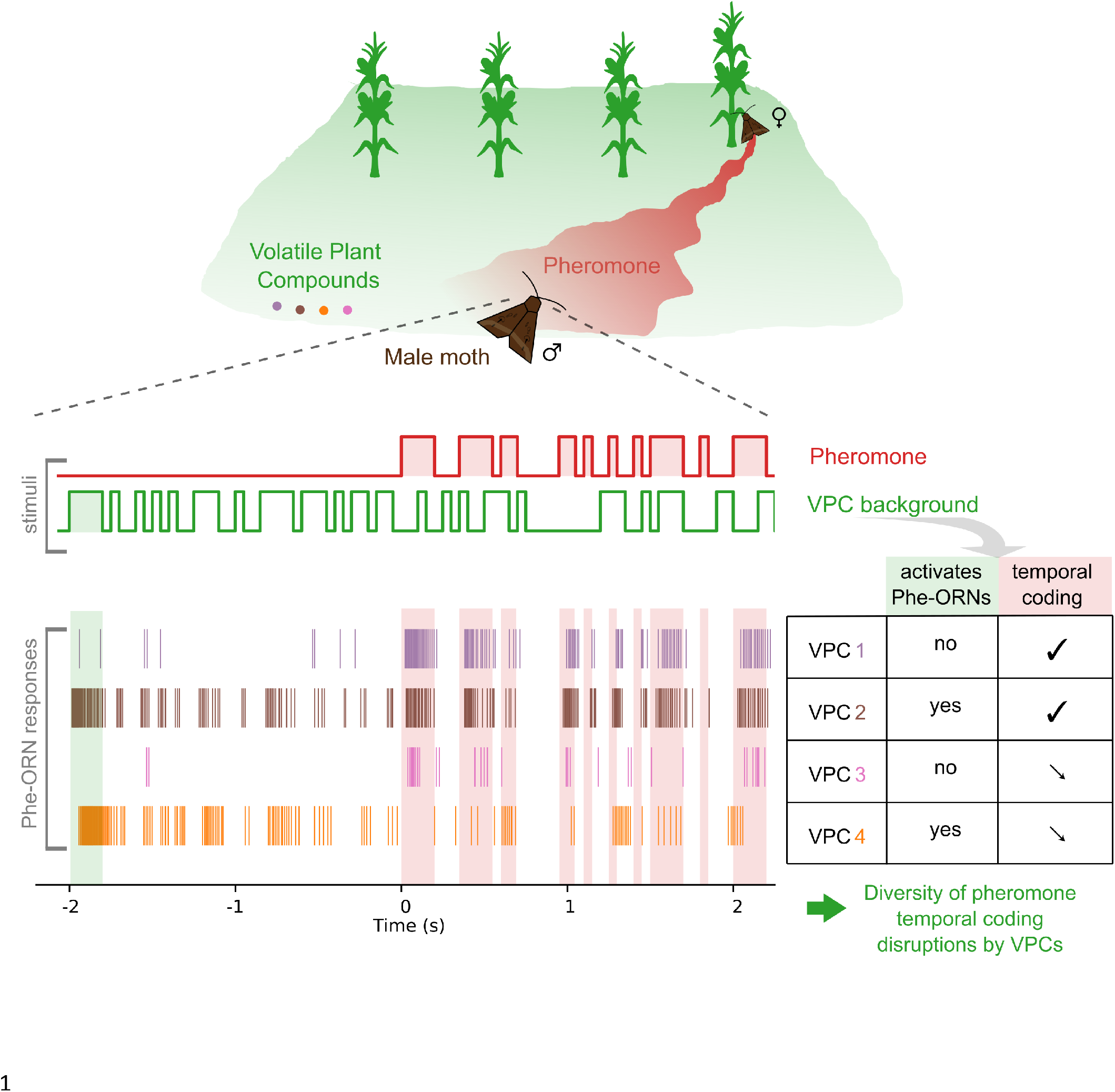

## Introduction

In many moths, males detect the sex pheromone blend components released by the female with their pheromone olfactory receptor neurons (Phe-ORNs), extract information about its intensity, quality, and temporal dynamics ^**1,2**^, and start a stereotyped orientation flight ^**3**^ towards the source. In natural environments, moths experience a very complex olfactory world whose composition changes in space and time ^**4–6**^. The surrounding vegetation releases a wide diversity of volatile plant compounds (VPCs), small secondary metabolites (<300 Da) ^**7–9**^. Despite the narrow tuning of Phe-ORNs to sex pheromones ^**10,11**^, several VPCs can activate Phe-ORNs, improving ^**12**^ or disrupting ^**13–19**^ pheromone detection. However, these studies relied on a single pheromone pulse or repeated square pheromone pulses of similar duration ^**20**^ delivered against constant odor backgrounds, conditions that remain far from the complex statistical structure of odor encounters in natural environments ^**21,22**^. In other insect models for olfaction, such as locusts or flies, the vast majority of studies presented ORNs at most with a single foreground odor against a constant odor background ^**23,24**^.

In natural environments, both VPCs and pheromones are mainly transported by three-dimensional air currents in a turbulent regime. Atmospheric turbulences create local vortices and instabilities, producing rapid fluctuations in odor concentration and wind direction, disrupting spatial gradients toward the source, and generating mixing between multiple odorants ^**21,25–32**^. As a result, insects encounter odor plumes characterized by high temporal variability and intermittency, rather than regular pulse trains. While the response dynamics ^**33–35**^ and efficiency ^**36–41**^ of the encoding of plume turbulent dynamics by insect ORNs were well-characterized in clean air, very little is known about how temporally structured olfactory backgrounds affect the neural encoding of the fine temporal structure of pheromone signals. Timing of pheromone encounters is thought to be crucial for olfactory navigation ^**42–44**^. Thus, any VPC-induced alteration of the temporal encoding of intermittent pheromone encounters would decrease the identification of navigation-relevant variables such as puff duration ^**38**^ or intermittency ^**45**^ among others, and ultimately lead to a decrease in the moth’s mating opportunities. To date, only a limited number of studies on locust projection neurons ^**36**^ and fruit fly ORNs ^**46**^, have investigated the neural response to turbulent odor *mixtures*, defined here as at least two different odorants presented simultaneously and delivered according to a fluctuating pattern. By contrast, how fluctuating olfactory backgrounds affect the encoding of intermittent pheromone signals at the level of moth Phe-ORNs remains unexplored. Importantly, the aforementioned studies on non-moth organisms investigated how relatively generalist olfactory systems encode odor mixtures composed of signals with comparable functional relevance ^**36**^, and did not specifically investigate background-induced modulations of the encoding of a behaviorally critical signal. On the contrary, the moth pheromone system is highly specialized ^**10**^ to track a single biologically relevant signal - the sex pheromone - embedded in ecologically relevant olfactory noise such as crop field VPCs that are known to modulate Phe-ORN gain ^**14**^.

Here, we stimulated Phe-ORNs of a noctuid moth, *Agrotis ipsilon*, with a sex pheromone delivered as a constant or a turbulent plume-like flickering stimulus consisting of pulses with variable durations and inter-pulse intervals, and in the presence of a diversity of ecologically relevant crop field odor backgrounds. We tested two temporal regimes by delivering a constant VPC background, mimicking emission from a large and spatially homogeneous source or a close source, or a fluctuating VPC background, mimicking unpredictable turbulent mixing between pheromone and VPC plumes ^**47,48**^. We tested a range of VPCs and quantified how they altered multiple dimensions of neural coding, including mean firing rate, spike timing precision, trial-to-trial variability, and the dynamics of spike-frequency adaptation. We showed that high concentrations of linalool, Z-3-hexenyl acetate (Z3HA), and some of their chemical congeners strongly altered the precision, gain, and reliability of pheromone temporal coding. We uncovered an unexpected diversity of VPC-induced effects, including cases in which temporal coding was degraded without detectable activation of Phe-ORNs, suggesting non-trivial peripheral interactions. The alteration of pheromone coding by VPCs cannot be predicted solely based on their ability to activate Phe-ORNs. These findings raise questions concerning the olfactory transduction cascade and downstream processing, and shed light on linalyl acetate as a compound of choice from a biocontrol perspective.

## Results

### Some VPCs decrease the gain and the ability of Phe-ORNs to encode the temporal structure of the pheromone

We first assessed the performance of the Phe-ORNs to encode the pheromone fluctuating signal in the presence of various VPC backgrounds (Figure 1, raw raster plots are shown in Supplementary Figure 1) by computing a logistic regression-based coding performance η. A constant (Figure 2A1) or fluctuating (Figure 2A2) background of eucalyptol, hexenal, β-caryophyllene and α-pinene does not strongly impact coding accuracy. By contrast, both a constant (Figure 2A1) and a fluctuating (Figure 2A2) background of Z-3-hexenyl acetate (Z3HA) and linalool strongly impair coding accuracy, and this effect is dose-dependent (Supplementary Figure 2). As Z3HA and linalool disrupt coding performance, we tested chemical congeners of these molecules to assess the specificity of their effect. We found that a constant ( Figure 2A1) or intermittent (Figure 2A2) background of (i) citronellol, a structural isomer of linalool, (ii) linalyl acetate, the acetylated form of linalool as well as (iii) E-2 hexenyl acetate (E2HA), a Z3HA-derived ester, also impairs coding accuracy. VPCs that unambiguously disrupt the pheromone coding (E2HA, linalyl acetate, citronellol, Z3HA, linalool) will be hereinafter referred to as ‘active VPCs’. Interestingly, a background of β-caryophyllene decreases significantly the mean firing rate (Supplementary Figure 3) but does not disrupt temporal coding (Figure 2A), indicating that a decrease in Phe-ORN gain is not always associated with a decrease in temporal coding.

**Figure 1.**
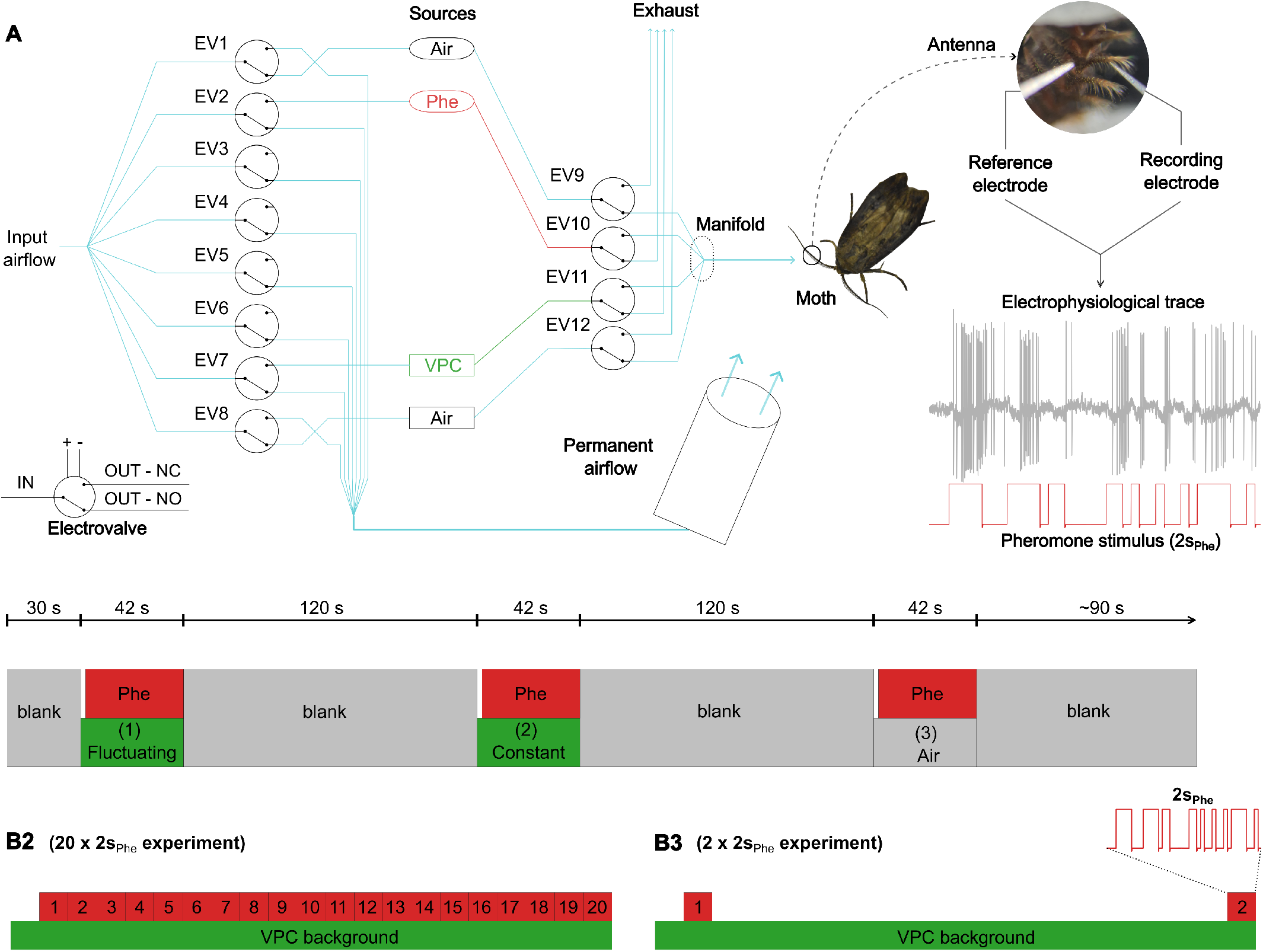
Experimental setup. **(A)**Experimental setup. A system of tubing and electrovalves enabled the delivery of a precisely timed mix of pheromone and a VPC while ensuring the delivery of a constant filtered and humidified air flow to a male moth antenna. The pheromone source is highlighted in red (“Phe”), and the VPC source is highlighted in green (“VPC”). Electrovalves are represented by circles with a switch inside. NC: Normally Closed. NO: Normally Open. The grey trace shows a raw electrophysiological trace of a representative Phe-ORN responding to the repeated 2-s pattern of pheromone (red) in the absence of background. **(B)**Stimulus sequences. (B1) Each sensillum received a 40-s pheromone stimulus (in red) three times, interspaced by 120-s blanks, yielding a total of a 8-min long recording. The 40-s pheromone stimulus was presented alone (protocol ‘air’), or in combination with a 42-s VPC stimulus delivered as a constant (protocol ‘constant’) or fluctuating (protocol ‘fluctuating’) background to the antenna. The order of presentation of the blocks (1) ‘fluctuating’, (2) ‘constant’ and (3) ‘air’ was pseudo-randomized to test each possible combination. (B2-B3) The 40-s pheromone stimulus consisted of a 2-s pheromone sequence (2s_Phe_, shown in the bottom right corner) repeated either 20 times (B2, protocol 20×2s_Phe_) or twice (B3, protocol 2×2s_Phe_).

**Figure 2.**
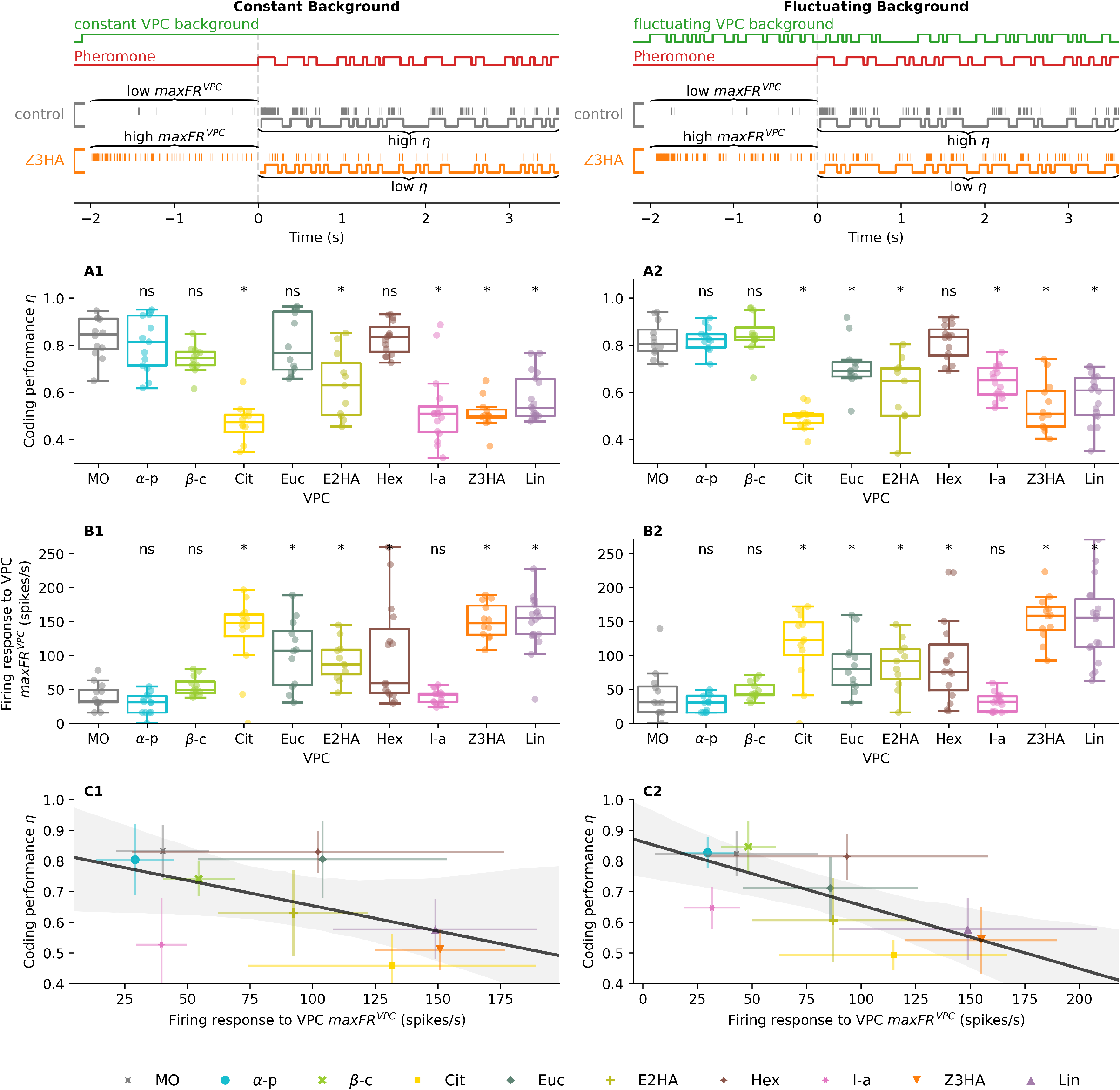
Correlation between the coding of the temporal structure of the pheromone in the presence of a plant volatile background, and the neural response to the plant volatile itself. In addition to the pheromone stimulus (shown in red), a plant volatile (shown in green) was applied either as a constant background (Left column, A1-B1-C1) or as a fluctuating background (Right column, A2-B2-C2). We computed the firing response to VPC (in the time window before pheromone onset, between -2 and 0 seconds) and the (de)coding performance, *i*.*e*. how well the original stimulus can be retrieved from the spike train, in the time window between 2 and 40 seconds. For illustration, we computed and showed the reconstructed stimulus between 0 and 2 seconds. **(A)**Coding performance (AUC score η) as a function of the tested VPCs. **(B)**Maximum firing rate during a 2-s window before the pheromone onset (maxFR_VPC_) as a function of the tested VPCs. Planned comparisons to mineral oil (MO), ns=non-significant, *: p-value <0.05. **(C)**Linear regression between the response to a VPC (maximum firing rate during a 2-s window before the pheromone onset) and η. The fitted regression line is shown in black, and the shaded area indicates the associated 95% confidence interval.

We observed that some VPCs significantly activate Phe-ORNs (Figure 2B, Supplementary Figure 3A) and decrease their firing response to the pheromone pulses: eucalyptol, citronellol, linalool, hexenal, Z3HA and E2HA. Thus, we investigated whether a decrease in the coding performance could be explained by a decrease in the Phe-ORN gain. We found a significant correlation between the coding performance η in the presence of a VPC background and the Phe-ORN response to this VPC (R^2^_constant_=0.34, Figure 2C1 and R^2^_fluctuating_=0.51, Figure 2C2, p<0.05 for both backgrounds), but with notable deviations. Intriguingly, linalyl acetate does not activate Phe-ORNs but significantly decreases the coding performance η (Figure 2A-B, Supplementary Figure 1, linalyl acetate: standardized residuals = -1.850, Cook’s distance = 0.524). On the contrary, eucalyptol and hexenal moderately activate Phe-ORNs but do not significantly impact η (Figure 2A-B, Supplementary Figure 1, hexenal: standardized residuals= 1.354, Cook’s distance =0.1; eucalyptol: standardized residuals= 1.363, Cook’s distance =0.1).

To complement these results, we assessed the coding efficiency by comparing various metrics: an estimate of the mutual information (Supplementary Figure 4A-B), the Jensen-Shannon distance between the spike train in the presence or absence of a VPC (Supplementary Figure 4C-D), or a proxy of the encoding performance, i.e., predicting the firing rate from the stimulus, with linear-nonlinear models (Supplementary Figure 5, Supplementary Figure 6). Overall, these results are consistent with the effects observed with the logistic regression index η. A short summary of the effects of VPCs can be found in Table 1.

**Table 1:** Summary of the effects of the tested VPCs. 0: no effect, +: increase, -: decrease, x: no data.

| VPC | Background | mean firing rate | max firing rate during VPC presentation | performance $\eta$ | encoding dynamics | contrast |
| --- | --- | --- | --- | --- | --- | --- |
|  |  | Supplementary Figure 3 | Figure 2 | Figure 2 | Supplementary Figure 5-6 | Supplementary Figure 7 |
| control (MO) | constant | 0 | 0 | 0 | 0 | 0 |
|  | fluctuating | 0 | 0 | 0 | 0 | 0 |
| Linalool (Lin) | constant | - | + | + | + | + |
|  | fluctuating | - | + | + | + | - |
| (-)-α-pinene (α-p) | constant | 0 | 0 | 0 | 0 | - |
|  | fluctuating | 0 | 0 | 0 | 0 | - |
| Eucalyptol (Euc) | constant | 0 | + | 0 | 0 | 0 |
|  | fluctuating | 0 | + | 0 (-) | 0 | 0 |
| β-caryophyllene (β-c) | constant | - | 0 | 0 (-) | 0 | - |
|  | fluctuating | - | 0 | 0 | 0 | - |
| Z-3-hexenyl acetate (Z3HA) | constant | - | + | + | + | + |
|  | fluctuating | - | + | + | + | - |
| E-2-hexenyl acetate(E2HA) | constant | - | + | + | + | + |
|  | fluctuating | - | + | + | + | - |
| Linalyl acetate (l-a) | constant | - | + | + | + | + |
|  | fluctuating | - | + | + | + | - |
| Citronellol (Cit) | constant | - | + | + | + | x |
|  | fluctuating | - | + | + | + | x |
| E-2-hexen-1-al (Hex) | constant | 0 | + | 0 | 0 | x |
|  | fluctuating | 0 | + | 0 | 0 | x |

### Pheromone stimulus subregions are differentially affected by a VPC background

We investigated which sub-regions of the pheromone stimulus (blanks and puffs) are most affected in their encoding by the presence of VPCs. We analyzed the logistic regression models’ accuracy of their predictions for each individual bin pooled for all repetitions (Figure 3, Supplementary Table 2). Overall, we did not find notable differences between a constant and a fluctuating background, but it should be noted that for citronellol, Z3HA, linalool, and E2HA, the response gain is higher in a fluctuating than in a constant background (Supplementary Figure 3B-C). For mineral oil (control) and non-active VPCs, the highest prediction errors relate to the end of long puffs, the short puffs, and the short blanks. For most active VPCs, VPC-induced spike-frequency adaptation produces sparse spike trains, preventing the logistic regression model from accurately predicting pheromone pulses and accounting for the higher error rates observed for these pulses. Interestingly, for linalyl acetate, prediction errors are distributed throughout the stimulus presentation (Figure 3), meaning that both puffs and blanks are badly encoded and that Phe-ORNs, rather than being completely in an adaptation state, completely lose track of the pheromone stimulus temporal structure.

**Figure 3.**
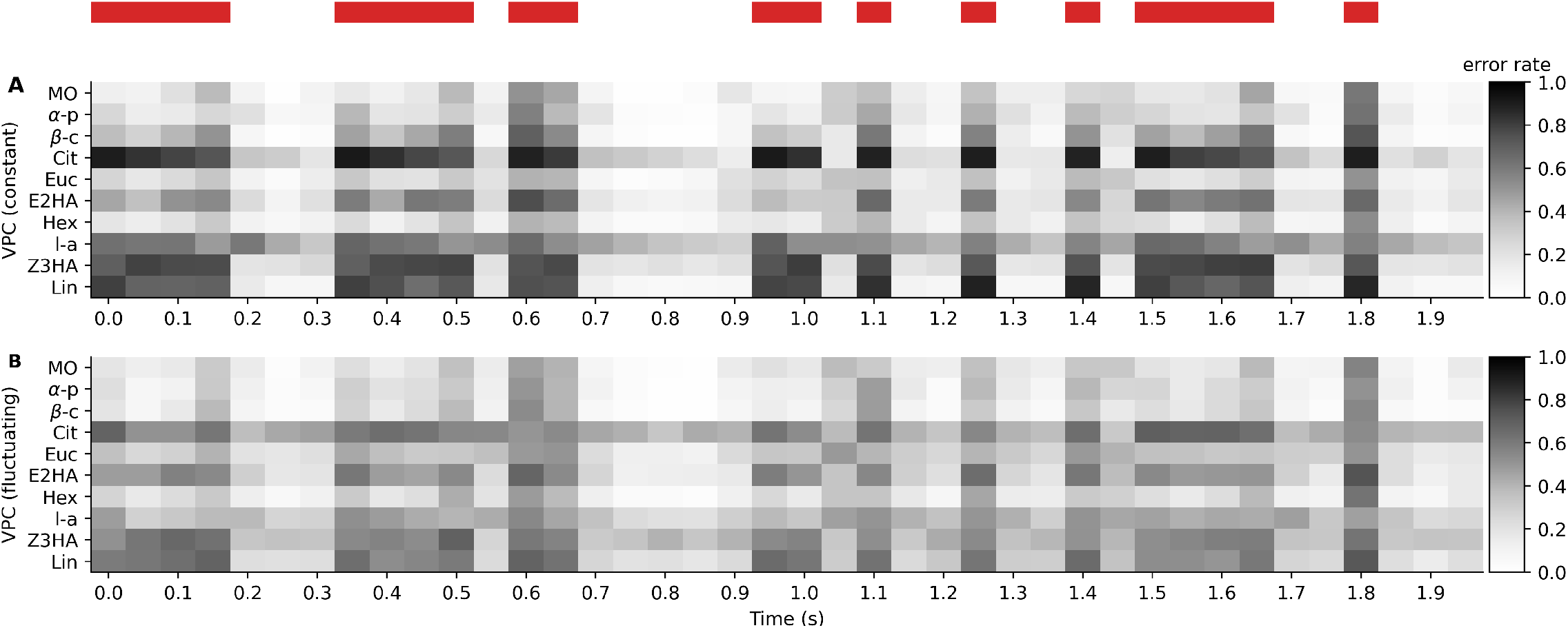
Neuronal coding of various time sub-regions of the 2-s pheromone stimulus. **(A-B)** Heatmap of the logistic model prediction mean error rate (greyscale colormap, with darker shades indicating higher error rates) for different VPCs (y-axis) as a function of time, in presence of a constant (A) VPC background or a fluctuating (B) VPC background. The 2-s pheromone stimulus is shown by red rectangles above the plots.

### Some VPCs alter the reproducibility of the response

We repeated 20 times the same 2-s sequence of pheromone stimuli to investigate whether a VPC background would alter the reproducibility of the response. Qualitatively, the jitter in the spike times appears lower in the presence of a control background (Figure 4A1) than in the presence of a Z3HA background (Figure 4A2). To quantify this jitter, we first performed a phase-locking analysis and compared the mean vector across VPCs, delivered as a constant ( Figure 4B1) or fluctuating background (Figure 4B2). For a constant or fluctuating background of citronellol, E2H, Z3HA, linalool and linalyl acetate, the mean vector is higher, indicating a higher variability between the spike trains associated with each repetition of the 2-s pheromone sequence (Figure 4). To confirm these results, we performed a multidimensional scaling analysis based on all possible pairwise distances between spike trains for each 2-s repetition (Figure 4C-D). Qualitatively, the dispersion ellipses are qualitatively similar for the control background (Figure 4C1), whereas they differ for the Z3HA background (Figure 4C2). This suggests that in the presence of a Z3HA background, the Phe-ORN response is less consistent across repetitions, indicating a higher trial-to-trial variability. We quantitatively compared the mean spike train distances for each tested VPC delivered as a constant (Figure 4D1) or fluctuating background (Figure 4D2). The mean distances are significantly higher in the presence of a citronellol, E2HA, Z3HA, linalool and linalyl acetate background, indicating an increased trial-to-trial variability consistent with the phase-locking analysis.

**Figure 4.**
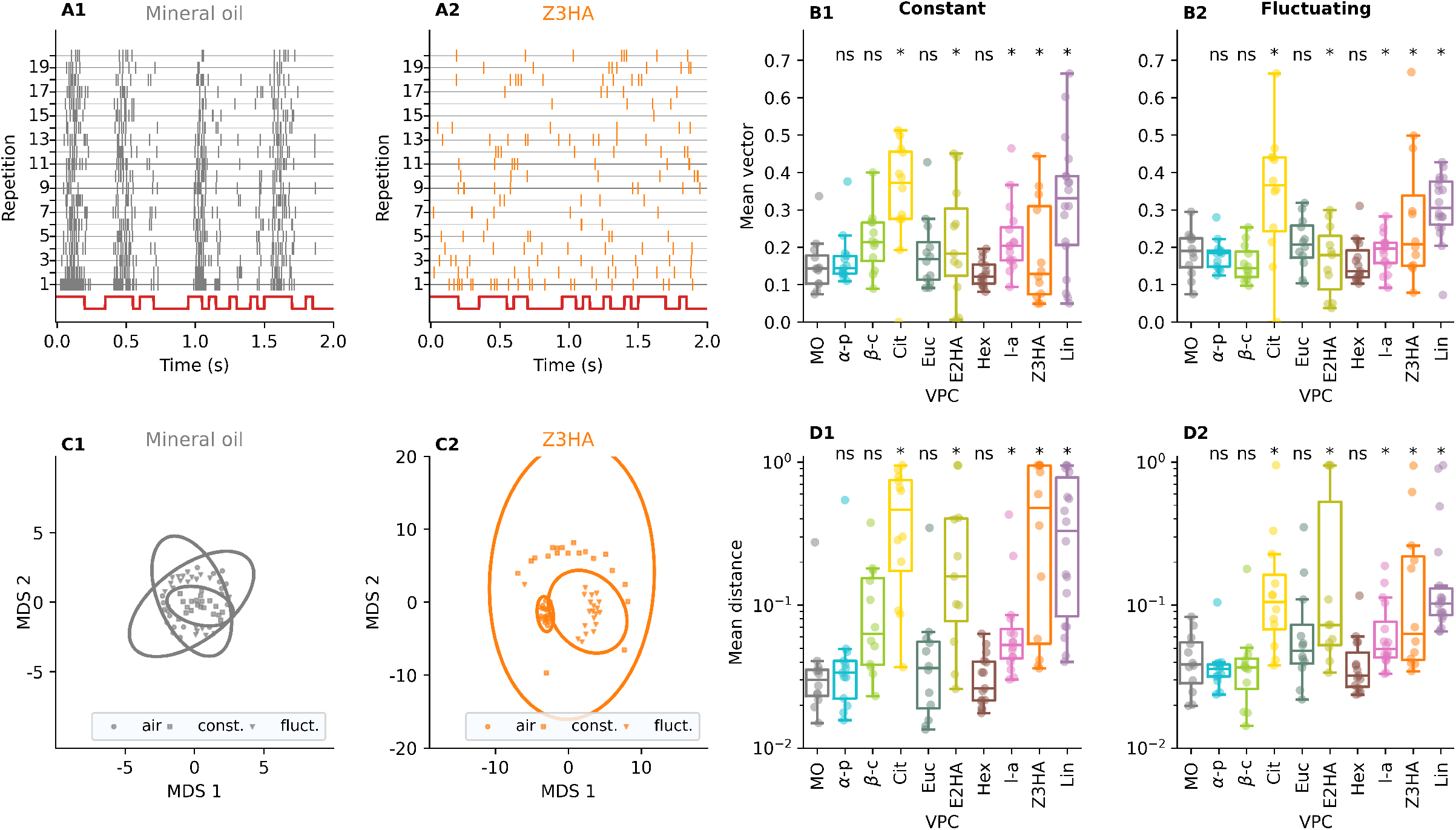
Some VPCs decrease the response reproducibility and increase the response jitter. (A-B) Phase-locking analysis. **(A)**Raster plot of a representative Phe-ORN in the presence of a fluctuating control (A1, grey) or Z3HA (A2, orange) background, where each row indicates a response to the 2-s repetition (total: 20 repetitions). **(B)**Boxplot of the mean vector strength for each tested VPC applied as a constant (B1) or fluctuating (B2) background. Higher vector strength values indicate a higher trial-to-trial variability. Planned comparisons to mineral oil (MO), ns=non-significant, *: p-value <0.05. **(C-D)** Multidimensional scaling (MDS) analysis based on the pairwise spike train distances. **(C)**MDS space and dispersion ellipse computed for a representative Phe-ORN in the presence of a fluctuating mineral oil background (C1, grey), and another representative Phe-ORN in the presence of a Z3HA (C2, orange) background. For a given recorded neuron, in which three distinct stimulation sequences were tested, circles indicate the 20 repetitions in the control background, squares indicate the 20 repetitions in the presence of a constant background, and triangles indicate the 20 repetitions in the presence of a fluctuating background. **(D)**Boxplot of the mean pairwise spike train distance for each VPC applied either as a constant (D1) or fluctuating (D2) background. Higher mean distances indicate a higher trial-to-trial variability. Planned comparisons to MO, ns=non-significant, *: p-value <0.05.

### VPC-induced effects on pheromone intermittency encoding in a protocol that reduces pheromone-induced adaptation

In the 20×2s_Phe_ protocol, the Phe-ORN response to the 20^th^ pulse (L-20p, last presentation - 20×2s_Phe_ protocol) is affected by the effects of adaptation to the preceding 19 pheromone sequences and the adaptation to the VPC background (Figure 1B2). We designed a protocol comprising only two presentations of the 2-s pheromone sequence separated by a 36-s pheromone-free period, where only an odor background is delivered to the antenna, to reduce the effect of pheromone adaptation (Figure 1B3). The first and last presentations were termed F-2p and L-2p respectively (first/last presentation – 2×2s_Phe_ protocol, Figure 5). We compared Phe-ORNs responses before and after a 36-s pheromone-free period. We investigated whether (1-2) the temporal coding performance and the gain to L-2p in the control and VPC background differ, (3) the responses to F-2p and L-2p stay similar. This protocol also allowed us to investigate putative late effects, i.e., whether the cumulative effect of a VPC over time can impair intermittency encoding.

**Figure 5.**
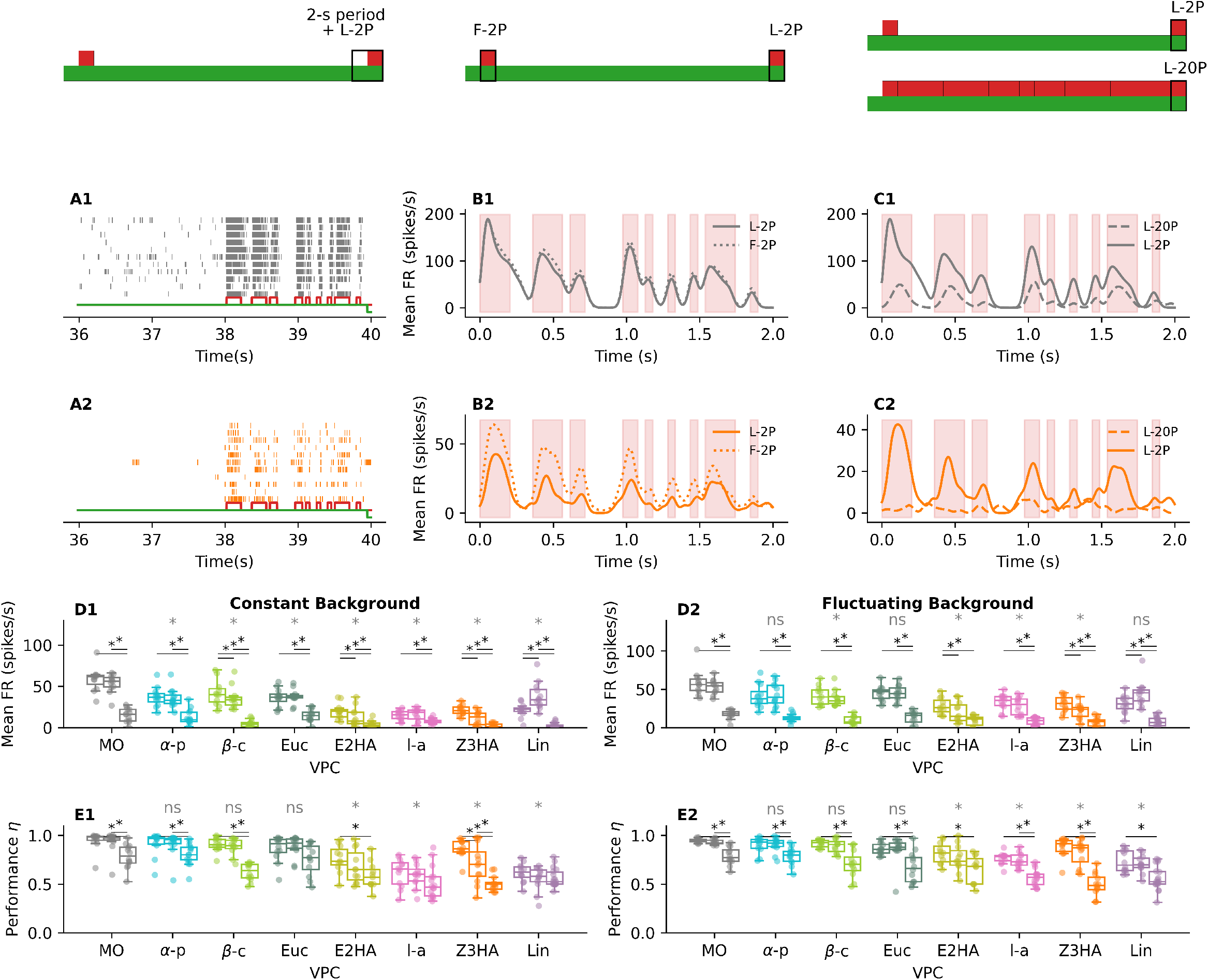
A 36-s presentation of an odor background without pheromone allows Phe-ORNs to recover from pheromone-induced adaptation under a mineral oil background, but not in the presence of a VPC background. **(A)**Raster plots showing the response of all recorded Phe-ORNs to the F-2p (Protocol ‘2×2s_Phe_’, time window between 36 and 40 s) in the presence of a mineral oil (A1, grey) or Z3HA (A2, orange) constant background. Note that, without any pheromone stimulation, a constant Z3HA background almost suppresses the firing activity of Phe-ORNs. The VPC stimulus is shown in green and the pheromone stimulus in red. **(B)**Mean firing rate (spikes/s) averaged over all recorded Phe-ORNs during the F-2p (dashed lines, Protocol ‘2×2s_Phe_’) and the L-2p (solid lines, Protocol ‘2×2s_Phe_’) stimulation in the presence of a mineral oil (B1, grey) or Z3HA (B2, orange) constant background. The light red rectangles show the pheromone stimulus. Note that, for mineral oil (B1), the two curves are superimposed, indicating that the Phe-ORNs recovered from pheromone-induced adaptation. **(C)**Mean firing rate (spikes/s) averaged over all recorded Phe-ORNs during the L-20p (dashed lines, Protocol ‘20×2s_Phe_’) and the L-2p (solid lines, Protocol ‘2×2s_Phe_’) stimulation in the presence of a mineral oil (C1, grey) or Z3HA (C2, orange) constant background. The light red rectangles show the pheromone stimulus. Note that, for mineral oil (C1), the response to L-2p is consistently higher than the response to L-20p. **(D)**Boxplot of the mean firing rate during the studied 2-s stimulation (for a given VPC, the grouped boxplots indicate, from left to right, F-2p, L-2p or L-20p) as a function of the tested VPC delivered as a constant (D1) or fluctuating (D2) background **(E)**Boxplot of the coding performance during the studied 2-s stimulation (for a given VPC, the grouped boxplots indicate, from left to right, F-2p, L-2p or L-20p) as a function of the tested VPC delivered as a constant (E1) or fluctuating (E2) background For (D) and (E), the black horizontal lines and stars indicate planned comparisons between F-2p and L-2p, L-2p and L-20p, and F-2p and L-20p per VPC. The grey stars indicate planned comparisons between L-2p in the presence of MO versus in the presence of a VPC background. (ns=non-significant, *: p-value <0.05)

We observed that a constant background of Z3HA or linalool almost suppressed the firing activity during the 36-s pheromone-free period (Figure 5A, interval between 36 s and 38 s). Despite this suppression, some Phe-ORNs responded to the pheromone pulses (a few spikes between 38 s and 40 s in Figure 5A, non-zero mean firing rate in Figure 5B-C-D), indicating that these backgrounds do not fully prevent pheromone detection. We quantified these observations by comparing the response *saliency*, defined as the normalized difference between the maximum firing rate before and during L-2p stimulation, across tested VPCs. Continuous active VPC backgrounds increased saliency by suppressing baseline activity (Supplementary Figure 7A1), whereas fluctuating active VPC backgrounds decreased saliency (Supplementary Figure 7A2), and both backgrounds decreased coding performance in comparison to the control background (Figure 5E). A constant background of VPCs previously identified as non-active (α-pinene) slightly decreased response saliency without altering coding performance. Overall, our results indicate that pheromone intermittency encoding cannot be inferred from response saliency alone. VPC backgrounds may transform pheromone encoding rather than simply abolishing pheromone responsiveness.

Moreover, in the presence of a constant control background, responses to F-2p and L-2p were qualitatively indistinguishable (Figure 5A1-B1), while responses to L-2p and L-20p differed in both their intensity and shape ( Figure 5C1). This indicates that Phe-ORNs recovered from adaptation after the 36-s presentation of the control background alone. In contrast, under a Z3HA constant background, responses to F-2p were higher than responses to L-2p (Figure 5A2-B2). It should also be noted that the response to L-2p more closely resembles the pheromone stimulus than the response to L-20p (Figure 5C2). We quantified these observations. First, the mean firing rate was consistently higher in response to L-2p than to L-20p for all VPCs and background conditions (constant: Figure 5D1, intermittent: Figure 5D2). For Z3HA, linalool, E2HA and β-caryophyllene, the mean firing rate significantly differs between F-2p and L-2p, indicating that such backgrounds prevented Phe-ORNs from recovering from adaptation. For linalyl acetate, the mean firing rate may be too low in response to F-2p to see any significant differences with L-2p. We also observed that, overall, the control background elicited higher responses to L-2p than VPC backgrounds. Second, in a constant background (Figure 5E1), the performance η differs between F-2p and L-2p only for Z3HA, and the performance worsens when compared to L-20p. This may indicate that for other active VPCs, the performance is already low for F-2p, resulting in an absence of differences with L-2p. In a fluctuating background (Figure 5E2), the coding performance was comparable between F-2p and L-2p for all VPCs, indicating that such a background did not degrade temporal coding. Except for E2HA and linalool, the performance was better for L-2p than for L-20p. When comparing L-2p across all tested VPCs, we observed that both coding performance (Figure 5E) and distance between F-2p and L-2p (Supplementary Figure 7B) were lower in the presence of mineral oil than in the presence of Z3HA, linalool, E2HA, linalyl acetate, and sometimes eucalyptol. This indicates an increased response variability and decreased coding performance in the presence of these VPCs. The mean firing rate was comparable between linalool and, e.g., α-pinene, but the coding performance is lower for linalool, suggesting that the response gain is not always an indicator of the quality of the temporal coding.

Altogether, these results demonstrate that prolonged VPC exposure does not silence pheromone-ORNs but reshapes the pheromone encoding, and can disrupt temporal coding without changing the response gain.

### Encoding of mixed turbulent odors

We have not found meaningful differences between a constant and a fluctuating VPC background (Figure 2). However, the reduced performance may be caused by different mechanisms. A constant background of linalool or Z3HA strongly reduces the gain (Supplementary Figure 3A1), dropping the firing rate to zero, and would prevent any spikes in response to the pheromone. By contrast, in a fluctuating background of such VPCs, spikes elicited in response to intermittent pulses of VPCs would interfere with coding of intermittent pulses of pheromone (Figure 6A).

**Figure 6.**
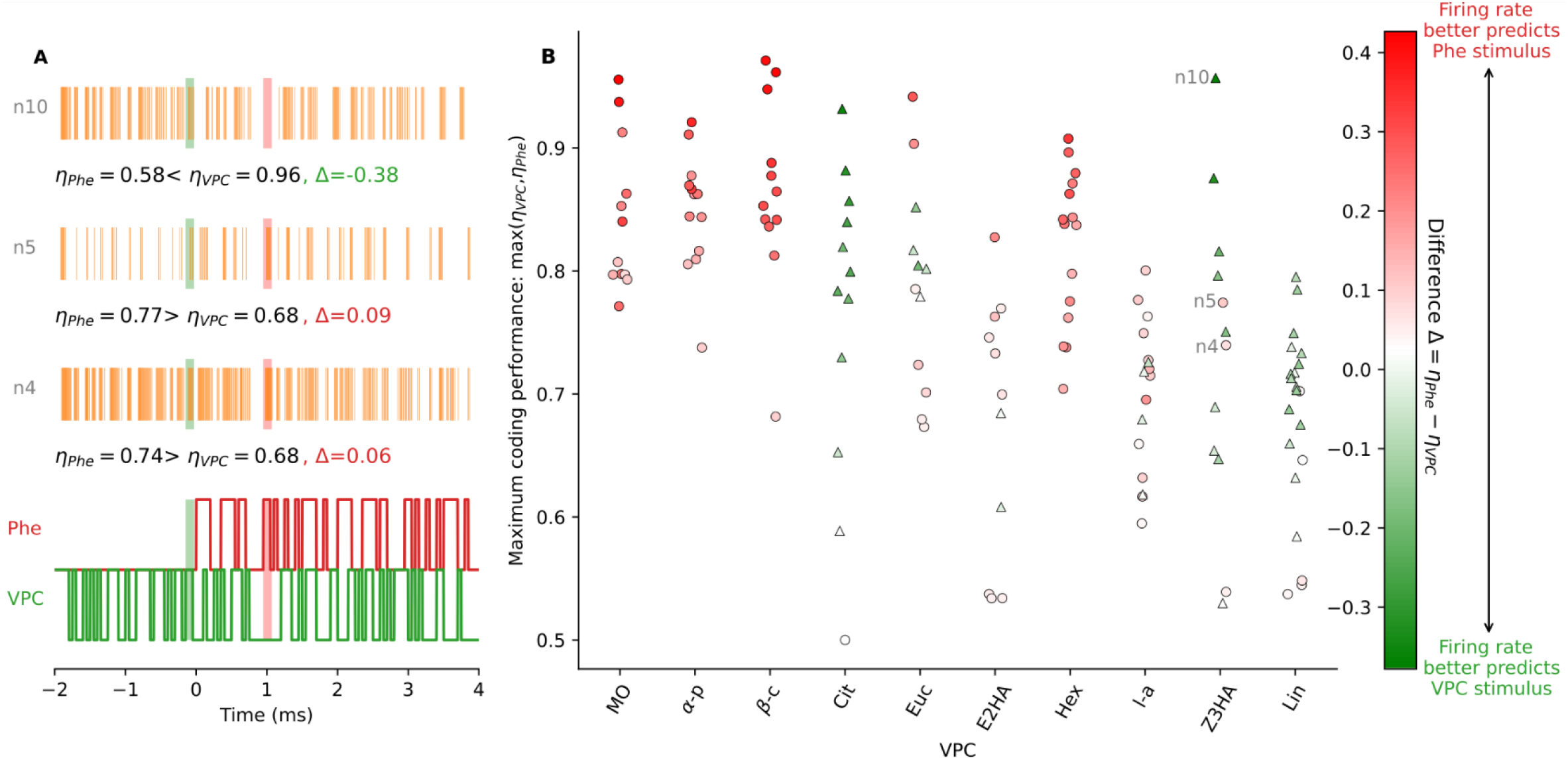
Phe-ORNs responses carry information about both the VPC and pheromone stimuli. **(A)**Raster plot (orange) of the response of some Phe-ORNs to a fluctuating Z3HA background (green trace) superimposed to a fluctuating pheromone stimulus (red trace). The green rectangle highlights a region where there is only a VPC pulse without pheromone (“VPC alone” pulse), whereas the red rectangle highlights a region where there is a pheromone pulse without VPC (“Phe alone” pulse). Note that the neuron n10 does not fire spikes in response to the “Phe alone” pulse, whereas the neurons n4 and n5 fire spikes in response to both the “Phe alone” and the “VPC alone” pulses. **(B)**Some responses cannot be explained solely by the pheromone stimulus alone. The logistic regression was performed on the pheromone values Phe(t), yielding the coding performance score η_Phe_, or the VPC stimulus values VPC(t), yielding the coding performance score η_VPC_. Each dot (square or disc) represents a single Phe-ORN. If η is greater when using the pheromone stimulus values as the predictor variable (η_Phe_ <η_VPC_), the dot was colored red and represented as a disc. Otherwise, the dot was colored green and represented as a triangle. The color intensity indicates the absolute value of the difference between η computed using either Phe(t) or VPC(t). Thus, a white dot indicates that no model performed really better than the other (η_Phe_ = η_VPC_).

Indeed, Phe-ORNs not only fire in response to pheromone pulses but also in response to linalool, Z3HA, and some of their derivatives. We investigated whether Phe-ORNs, when subjected to a mix of intermittent pheromone and VPC, would encode the pheromone stimulus, the VPC stimulus, or a combination of both (Figure 6A). To get a first qualitative overview, we computed the accuracy of a logistic regression model using either the pheromone stimulus or the VPC stimulus as the input and selected the model with the highest accuracy (Figure 6B). For mineral oil and non-active VPCs, the pheromone stimulus is consistently better at predicting the spike trains than the VPC stimulus (red dots, scores >0.8). For active VPCs, in most cases neither stimulus is really better than the other in predicting the spike trains (white dots, scores ∼0.5-0.6). We noticed that in a few cases, the spike trains are better predicted by the VPC stimulus (green dots, scores >0.8). This indicates that, in the presence of an active VPC fluctuating background, the spike trains cannot be predicted using the pheromone stimulus only. However, this approach does not allow us to directly determine whether Phe-ORNs encode one or both stimuli. Indeed, the raster plot shown in Figure 6A qualitatively highlights that some neurons can respond to both the pheromone and the VPC stimuli.

Then, we fitted a generalized linear mixed model (GLMM) to model the Phe-ORN response as a function of the pheromone and VPC stimuli. Based on a comparison between an additive and an interactive model, we also included the interaction between the VPC and the pheromone to unravel potential non-linear effects in the coding (Supplementary Table 3). For instance, for Z3HA, both pheromone (β= 1.045, SE = 0.058, p<0.001) and Z3HA (β=2.180, SE = 0.051, p<0.001) significantly increased the neural response. We also observed a strong negative interaction between pheromones and VPC (β= –1.134, SE = 0.063, p<0.001), suggesting that the simultaneous presence of VPC significantly attenuated the neuronal response to the pheromone. This indicates that Z3HA exerts a suppressive modulatory effect on pheromone-evoked activity, potentially reflecting early-stage mixture nonlinear interactions in olfactory coding. A complete summary of GLMM coefficients across all VPCs is provided in Supplementary Table 3. Notably, the majority of VPCs exhibited similarly significant negative interactions with the pheromone, reinforcing the idea that olfactory encoding of mixtures is shaped by odor-specific, non-additive integration dynamics.

## Discussion

Understanding how sensory neurons extract biologically relevant signals from noisy environments is a central question in sensory neurobiology. Previous studies in moth olfaction focused either on how continuous VPC backgrounds modulate the detection of simple pheromone pulses ^13–15,19,20^, or explored how Phe-ORNs track complex temporal structures in synthetic pheromone plumes in the absence of background odors ^39,40^. These two lines of research—pheromonal signal intermittent structure and background interference—remained largely disconnected. However, analyzing how VPCs influence the coding of the temporal dynamics of pheromonal stimuli is crucial, as the encoding of pheromone pulse duration is highly ecologically relevant ^38^—especially for plume-tracking behaviors (Balkovsky et Shraiman 2002; Demir et al. 2020). In our study, we bridged this gap by evaluating how VPC backgrounds affect the encoding of the temporal dynamics of the pheromone stimulus. VPC stimuli were delivered either as a constant background, mimicking emission from a large and spatially homogeneous plant source ^47^, or as a temporally fluctuating background ^48^, mimicking turbulent mixing between plant volatiles and pheromone plumes typically encountered in natural odor landscapes. We systematically quantified how VPCs, 6 tested in ^14^ and 3 new ones (citronellol, linalyl acetate, and E2HA), impact the neural representation of pheromone intermittency in the noctuid moth *Agrotis ipsilon* by computing multiple complementary metrics: gain, latency, firing precision and signal reliability, among others, whose dependence to the mean firing rate differs.

### Implications for moth navigation

Our results indicate that both a constant and fluctuating VPC background can impair the coding precision of pheromone temporal structures at the peripheral level. Some VPCs reshape pheromone encoding through non-additive interactions at the ORN level rather than simply adding noise and compress the dynamic range available for pheromone-evoked firing. As a consequence, the temporal structure of the plume that is available to downstream circuits is already distorted, disrupting the identification of puff onset and duration, plume intermittency, or local gradients, which are key for successful olfactory navigation. This should result in a decrease in the coding precision of antennal lobes neurons ^14,15,19,49^ and a disruption of olfactory navigation in wind tunnels ^15,20^. Moreover, theoretical studies showed that alterations of the sensors’ coding precision indeed reduce the available spatial information that can be used for navigation and reduce the probability of an agent finding the source ^50,51^. However, a disruption of coding at the peripheral level may not imply a disruption of olfactory-mediated behaviors. Despite a high dependency of the sensory input on background odors, mitigating mechanisms such as sensory averaging via temporal and spatial summation ^52^, followed by a dynamic reorganization of the activity pattern of higher-order neurons ^23,53^ via specific innervation patterns ^54^ may lead to background-independent odor recognition and potentially robust olfactory navigation. Moreover, in nature, the pheromone signal and VPC cues perceived by moths would be asynchronous, providing sufficient information to discriminate between the two sources ^32,55,56^. To assess whether moth tracking behavior is altered by the constant and fluctuating background we tested, a first step would consist in analyzing the tracking behavior of restrained moths on a spherical treadmill ^20^, and stimulating them with the same stimulus device we used for our electrophysiological experiments.

It should be noted that temporal coding remained unaffected at all lower VPC dilutions we tested, raising questions concerning the ecological relevance of the highest concentrations used in this study. However, VPC measurements are usually averaged over time and space (e.g. 8 ng/hour/plant, ^9^), and moths could indeed experience high local concentration peaks. Furthermore, with increasing temperatures, plants will also undergo more stress and release VPCs in higher quantities ^57–59^ resulting in a potential loss of mating opportunities ^60–62^.

### Some VPCs degrade the encoding of the pheromone temporal structure

VPCs do not affect Phe-ORNs equally: while some have minimal influence, others substantially degrade the encoding of the pheromone signal. Overall, there is a negative correlation between the intensity of the response induced by the VPCs and the pheromone coding efficiency in the presence of VPCs. The activation of Phe-ORNs by VPCs and the inhibitory effect of some VPCs are consistent with a previous study ^14^ which investigated the detection of a single pheromone pulse in the presence of similar VPCs. Whereas studies on VPCs ^19,37,63^ have focused only on saliency and response intensity, the biologically meaningful information is the capacity to encode both the onset and the duration of the pheromone pulses, and we pointed out that neither the response intensity nor the saliency is a good proxy. We found that a constant (but not fluctuating) VPC background can, to some extent, increase the response saliency to pheromone pulses by reducing the background activity, consistently with the aforementioned studies, but we highlighted that saliency does not reflect temporal coding performance. Indeed, a constant background of linalool, for instance, increases the saliency of the first pheromone pulse, but overall, the coding performance is far worse than the control background. In this study, we showed here that some VPCs degrade the ability of Phe-ORNs to track pheromone signal intermittency.

We initially expected a simple dichotomy between VPCs that activate Phe-ORNs and decrease temporal coding, and VPCs that do not activate VPCs and do not disrupt temporal coding. Interestingly, while the ability of the Phe-ORNs to encode the temporal structure of the pheromone is overall correlated with the response to the VPC alone, we observed two notable deviations from the regression line. Linalyl acetate, the acetate ester of linalool found in many flowers ^64^, does not activate Phe-ORNs but disturb coding performance. Conversely, hexenal and eucalyptol moderately activate Phe-ORNs but do not appear to degrade coding performance. β-caryophyllene reduces the mean firing rate of Phe-ORNs but it does not impair temporal coding. To sum up, the alteration of pheromone coding by VPCs cannot be predicted solely based on their ability to activate Phe-ORNs.

We expected that coding performance would be different under a constant background than a fluctuating background because their temporal statistics diverge—constant backgrounds are more predictable but exhibit fewer transient events ^65^. We found no clear differences in the raw coding performance scores between constant and fluctuating backgrounds. However, the mechanisms underlying the decrease in coding performance may differ. Our results suggest that a constant background of Z3HA or linalool induces strong adaptation and suppresses further responses to pheromone pulses, whereas a fluctuating Z3HA and linalool background interferes with pheromone pulse encoding. In other sensory systems, both constant and temporally fluctuating backgrounds are known to impair the encoding of superimposed signals, but they do so via partly distinct mechanisms: continuous backgrounds compress responses to additional stimuli, whereas fluctuating backgrounds distort the temporal structure even when mean firing rates are modestly reduced ^66–68^.

### Adaptation kinetics of Phe-ORNs in presence of a noisy background

Our results show that prolonged exposure to particular VPCs (Z3HA, linalool and some derivatives) produces a modulation of Phe-ORN responsiveness that is not simply explained by spike-frequency adaptation to repeated pheromone pulses. Although tonic firing is strongly reduced under continuous VPC backgrounds, pheromone pulses can still evoke spikes, albeit with altered latency and precision. This indicates that background VPCs neither saturate nor irreversibly block the transduction pathway, but instead alter the dynamics of pheromone coding, leading to degraded temporal representations. The pheromone receptor *Aips*OR3 appeared to be very specific to Z7-12:OAc and not to VPCs ^69^, indicating that the adaptation caused by the VPCs would not result from the binding of VPCs on *Aips*OR3. Although the mechanisms remain unclear, adaptation induced by VPCs is thought to be mediated via a putative unidentified VPC-sensitive receptor expressed in Phe-ORNs which would alter pheromone response dynamics by depolarizing Phe-ORNs.

We observed four distinct VPC effects on Phe-ORNs: (i) VPCs that do not elicit spikes and do not disrupt the pheromone coding, such as α-pinene (ii) VPCs that trigger a firing response and degrade pheromone response, such a Z3HA (iii) VPCs that elicit moderate spiking without substantially altering pheromone responses, such as eucalyptol, and (iv) VPC that does not elicit spiking activity yet still disrupt the pheromone coding (linalyl-acetate). Linalyl acetate and Z3HA likely alter temporal coding via different mechanisms: linalyl acetate might be a pheromone receptor antagonist, while Z3HA appears to directly evoke firing possibly by binding to a different, putative receptor expressed by Phe-ORNs ^69^. By contrast, although eucalyptol significantly activates Phe-ORNs, it did not degrade pulse tracking or temporal resolution of pheromone responses, suggesting that the mechanisms underlying tonic activation by eucalyptol and phasic pheromone encoding are at least partially segregated. These findings raise questions concerning the olfactory transduction cascade, and deeper investigations would require *in silico* docking approaches ^70^ and analyses of the moth pheromone tracking behavior ^19,20^ in the presence of such plant volatiles. Linalyl acetate, a putative Phe-ORN antagonist which has unexpected disruptive effects on pheromone encoding, is a candidate of choice for such analyses. Verifying its specificity of action toward the pheromone receptor of *A. ipsilon* will be the next step, and it may also hold significant potential from a biocontrol perspective ^71^.

In summary, we demonstrate that VPC backgrounds have a diversity of effects, indicating that the pheromone detection system must contend with multiple forms of background noise rather than a uniform disturbance. The tracking behavior should therefore be compared across different VPC backgrounds and with sources of different sizes to assess the behavioral relevance of our electrophysiological findings. These results are also relevant for the development of electronic noses ^72,73^, where the possible interferences are multiple. Noise mitigation in such systems may require a combination of techniques tailored to the unique features of the variety of background noises rather than a single solution.

## Materials and methods

### Animal care

Larvae of *Agrotis ipsilon* (Hufnagel, Lepidoptera: *Noctuidae*) were reared on an artificial diet ^74^. Males were separated from females at the pupal stage (the pupae have a sexual dimorphism) to prevent them from being exposed to the pheromone before the experiments. New adults were collected every day, kept under a 16h:8h light:dark inverted photoperiod at 20°C and fed *ad libitum* with a sucrose solution. Male were tested at 4 to 5 days old, when they have reached sexual maturity. All experiments were performed during the scotophase.

### Chemicals

For the VPCs, we chose linalool, α-pinene, Z-3-hexenyl acetate, E-2-hexenal, and eucalyptol because of their variety of physicochemical properties and their ecological relevance for a male moth seeking a female in a French agroecosystem ^75– 77^. As we observed an effect for linalool and Z-3-hexenyl acetate, we tested the response of Phe-ORNs to some chemical congeners of these active molecules: linalyl acetate, citronellol, and E-2-hexenyl acetate. Synthetic standards, CAS numbers, and their abbreviations are presented in Supplementary Table 1 and were acquired from Sigma Aldrich (Sigma Aldrich, Saint-Quentin Fallavier, France). We used the partition coefficients measured by Conchou and colleagues ^78^ to compensate for differences in volatility and to allow consistent comparisons between VPCs. All delivered VPC concentrations were defined relative to an arbitrary unit, where one unit is the molar concentration delivered on the antenna from a source containing 1% VPC in mineral oil. For each VPC, the appropriate concentration vol/vol to account for differences in volatility is presented in Supplementary Table 1. Mineral oil alone was tested as a control background. Odor sources were prepared each day, prior to any experiment, under an extractor hood located in a distinct room from the one used for electrophysiological recordings.

We used (Z)-7-dodecenyl acetate (Z7-12:OAc, Pherobank, Duurstede, Netherlands), one of the major components of the *A. ipsilon* sex pheromone blend as the pheromone stimulus, as used previously ^39,40^. Doses of 0.1 ng of Z7-12: OAc diluted in hexane (CAS 110-54-3, Carlo-Erba, Val-de-Reuil, France) were used in all experiments.

### Odor stimulus delivery

The experimental setup enabled the delivery of a precisely timed mix of pheromone and a plant volatile. It consisted (Figure 1A) of a set of three-way solenoid electrovalves (LHDA1233215H, The Lee Company, Voisins-le-Bretonneux, France) delivering both pheromone and VPC to a low dead-volume 4-way-manifold (MPP-8, Warner Instruments, Holliston, MA, USA) to ensure mixing. The air input consisted of a continuous flow of filtered (refillable hydrocarbon trap, Restek France, Lisses, France) and humidified air, limiting air contamination and preventing the antenna from drying out. This flow was divided into eight parallel streams thanks to an airflow divider (LFMX0510423B, The Lee Company, Voisins-le-Bretonneux, France), each directed to a solenoid valve (Figure 1A, EV1-EV8). PTFE tubing (1.32 mm ID, PTFE Tube Shop, Netherlands) connected the air input to the odor sources, the electrovalve, and the manifold.

The VPC sources consisted of 1 mL glass vials containing the VPC diluted in mineral oil, as described previously ^20^. They were kept at 23°C using a Thermo Mixer (ThermoMixer C, Eppendorf, France) to avoid temporal variations in headspace concentrations. Vials were sealed with a Teflon septum, and a combination of hypodermic needles and Teflon tubing connected the flow input to the odor vials, an electrovalve, and the manifold. The pheromone source was composed of a 3 cm-long capillary containing a filter paper impregnated with 0.1 ng of Z7-12Ac, a dose described as the active concentration ^19^, diluted in 1 µL of hexane. The pheromone was sent to the manifold and thus to the antenna only if the associated electrovalve was switched on (Figure 1A, EV10), and was otherwise sent to evacuation to limit room contamination. The valve delivering the pheromone (Figure 1A, EV9) was driven in antiphase with a valve delivering odor-free air, thereby ensuring a constant airflow. The same was true for the valve delivering the VPC (Figure 1A, EV11-EV12).

The outlet of the manifold was positioned close (<1 cm) to the recorded sensillum. Contaminated air was removed using an exhaust fan. Electrovalves were powered with a voltage generator (12V).

Pheromone and odor sources were changed between each recording. We ensured every day that the air flow at the distal end of the manifold in all possible different valve configurations used in our experiment was equal to 200 mL/min (± 10 mL/min) with an electronic flowmeter (GFM Pro Gas Flowmeter, ThermoFisher Scientific, France). Each day, or whenever a new VPC was tested, we decontaminated the tubing, the electrovalves, and the manifold at 80°C during 4 hours. The stimulus device was calibrated by recording the manifold output with a photoionization detector (Mini-PID, Aurora Scientific Inc., Canada), as described in previous studies ^20^.

### Single Sensillum Recordings

For electrophysiological recordings, we used tungsten electrodes (TW5-6, Science Products, Hofheim, Germany) sharpened electrolytically. Male moths were quickly anesthetized with CO_2_ and restrained in a styrofoam holder with their heads protruding. Once an antenna was immobilized with adhesive tape, the reference electrode was inserted in the flagellum, and the recording electrode was inserted at the base of a trichoid sensillum located on a ramus ^79^, using an NMM-25 micromanipulator (Narishige, London, UK). The signal was amplified (x1000) and band-pass filtered (0.1 - 10 kHz) using an EX1 amplifier with a 4002 headstage (Dagan, Minneapolis, USA). The signal was then digitized by a Digidata 1440A acquisition board (Molecular Devices, San Jose, USA) and sampled at 10 kHz under Clampex 10.3 (Molecular Devices, Sunnyvale, CA, USA). We recorded one sensillum per insect.

### Stimulation sequences

We recorded each Phe-ORN during an eight-minute stimulation consisting of the presentation of 3 distinct 40-seconds stimulations (‘fluctuating background’, ‘constant background’, and ‘no background’) separated by a 2-min blank (Figure 1B1).

Control of electrovalves was done by custom-made LabVIEW (National Instruments, Austin, USA) programs running on an NI-9472 module (National Instruments, Austin, USA). Stimulation sequences were defined in text files, where the state of each electrovalve was represented as a Boolean variable. The correlation time was 50 ms, indicating that the valve state was updated every 50 ms. For the generation of pheromone and VPC fluctuating sequences, the durations of whiffs and blanks were drawn from a truncated power-law distribution (∼t^-3/2^) to mimic the turbulent dynamics of an odor plume, following the model of Celani et al. ^21^. The probability density function of whiffs and blank duration (t) was given by:

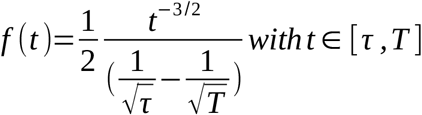

where τ and T are lower and upper cutoffs determined by the mean wind velocity U, the wind fluctuation velocity δU, the pheromone source size a and the intermittency factor χ.

For the pheromone, a 2-s sequence composed of nine pheromone puffs of random durations, interspaced by blanks of random durations (see red trace in Figure 1A-Figure 1B3), was generated as described above. In a first experiment (20×2s_Phe_ experiment), this 2-s pheromone sequence was repeated 20 times (Figure 1B2). In a second experiment (2×2s_Phe_ experiment), we changed the experimental protocol to investigate spike-frequency adaptation, and we only kept the first (between 0 and 2 s) and the last (between 38 and 40 s) 2-s pheromone sequence presentations (Figure 1B3). In such a protocol, the first and last presentations will be hereinafter referred to as ‘F-2p’ (First presentation – 2×2s_Phe_ protocol) and ‘L-2p’ (Last presentation – 2×2s_Phe_ protocol) respectively.

In the meanwhile, the VPC background was either (1) delivered as a 42-s fluctuating pattern (protocol ‘fluctuating’ in Figure 1B), (2) delivered continuously (protocol ‘constant’ in Figure 1B), or (3) absent (control stimuli, protocol ‘air’ in Figure 1B). The background onset started 2 ms before the pheromone stimulation to assess the response of Phe-ORNs to individual VPCs (Figure 1B1, Supplementary Figure 1); thus, the onset of VPC stimulus is defined at t=-2 s. The 3 distinct stimulations — (1) pheromone + fluctuating background, (2) pheromone + constant background and (3) pheromone + no background —were randomly presented to the moth. Therefore, there were 6 different recording sequences (3! = 6 permutations), and we tested each sequence at least twice. Thus, we recorded the response of the Phe-ORNs of at least 12 different moths per background (Supplementary Figure 1).

For each recorded neuron, the internal control consisted in testing the firing response to the pheromone without any VPC background (protocol ‘air’). Testing the effect of the solvent on each recording neuron was experimentally challenging and would require more electrovalves. We assessed the contribution of the solvent (mineral oil). No significant differences between the treatments ‘No background’ (protocol ‘air’) and ‘mineral oil background’ were found (Supplementary Figure 8A). Thus, downstream analyses assumed that solvent effects did not account for differences between the ‘No VPC’ and ‘VPC’ treatments, and we referred to experiments where a mineral background (solvent control) was applied and experiments where no odor background (protocol ‘air’, internal control) was applied as the ‘control background’ experiments.

## Data quantification

The custom-written Python (version 3.11, Python Software Foundation, https://www.python.org/) scripts used for the downstream analyses can be found in an online Github repository: https://github.com/pclemenc/FluctuatingVPC. We used diverse complementary metrics differing in their sensitivity to the mean Phe-ORN gain. The key question is whether Phe-ORNs encode the duration (onset and offset) of pheromone pulses, irrespective of the total number of spikes elicited.

### Spike timing extraction and computation of the firing rate

Spike timing extraction was performed with the Spike2 software (CED, Oxford, UK). Valve opening and closing times, as well as spike occurrence times were exported to a .txt file using a custom-written script as used previously ^80^. We estimated the firing rates by the kernel density estimation method ^81^. Each spike was substituted with a normal distribution probability distribution function with a mean at the spike time and a variance of 50 ms.

### Binary logistic regression

We used binary logistic regression as a proxy for the decoding performance: how a naïve classifier, based on the spike train, can deduce the original stimulus sequence. It is a proxy of the information that would be available for the downstream neuronal populations.

For the 20×2s_Phe_ experiment, we performed a logistic regression between the spike counts and the pheromone stimulus values s_Phe_(t). The spikes were binned in 50 ms intervals, as it is the correlation time of the valve state, yielding a total of 800 bins. In order to remove the short-term adaptation, we removed the first 2-s sequence for the subsequent analysis. Thus, there are 800-40=760 bins in total. We first performed a logistic regression by using 380 random bins as training data and the remaining 380 bins as test data on the spikes associated with the ‘air’ protocol. The training data allowed us to find the parameters of the logistic regression model, and this model was applied to the spikes associated with the constant and fluctuating background conditions for subsequent performance assessments. The performance was evaluated by the Area Under the Receiver Operating Characteristic Curve (AUC) metric, which equals 0.5 if the prediction is no better than a random prediction and 1.0 if the prediction is perfect. In order to take the delay between the neural response and the electrovalve onset into account, we time-shifted the bins by various delays δ (between 0 and 85 ms) and computed the associated AUC_δ_. We defined the temporal coding logistic-based performance η as the maximum computed AUC: η = argmax(AUC_δ_), _δ € [0,85]_.

For the 2×2s_Phe_ experiment, the same procedure was applied, but we kept the first 2-s trial, yielding a total of 80 bins. In parallel, we also divided the binned spikes associated with each protocol (air/absent, constant, fluctuating) into 380 train bins and 380 test bins, and thus performed three independent logistic regressions for each protocol and computed the best AUC values, named κ to avoid confusion with η values described above. This approach was less privileged because it does not use the absent background (control) protocol as explicitly as the first approach but led to similar results (Supplementary Figure 8B).

### Coding accuracy of different subregions of the stimulus

In order to understand which specific parts of the stimulus are particularly poorly encoded, we aligned the predictions of the logistic regression with the stimulus time course. Incorrect predictions (false positives and false negatives) were flagged as errors. We averaged the 20 repetitions of the 2-s pheromone stimulus to retrieve a mean error rate.

### Kernel estimate analysis

We fitted linear-nonlinear models to model the firing activity of each neuron in order to better characterize the Phe-ORNs’ temporal properties. However, the pheromone stimulus consists of the repetition of the same repeated 2-s sequence, and is thus not a perfect white-noise stimulus with the required diversity of possible whiff and blank durations. This approach complements the decoding performance metrics described above and will be interpreted only as a rough proxy of the encoding performance. We retrieved the linear kernels by following a method previously described ^36,39^. Let s_Phe_(t) be the stimulus. Briefly, the neuron firing rate r(t) can be written as a combination of a linear kernel K and a nonlinear function F: r(t)=F(**∫** K(t-u).s_Phe_(u).du). The linear kernel K was estimated using an elastic net regression (function sklearn.linear_model.ElasticNet in Python) between the firing rate and the time shifted stimulus matrix in a short time window (-1 to 1 s). A nonlinear function was then determined to compensate for the amplitude of the filter by fitting a threshold rectifier (ReLU) between the predicted firing rate by the linear filter, and the measured firing rate with the scipy.optimize.curvefit function in Python. The nonlinear function requires two parameters (gain and threshold).

In order to analyze the diversity in the kernel shapes, which characterizes how a neuron encodes a fluctuating stimulus, we subjected them to a principal component analysis. We extracted the two eigenvectors P_1_(t) and P_2_(t) with the highest eigenvalue. The kernels K were normalized in amplitude so that each filter corresponds to one dot with coordinates x=**∫**K(t).P_1_(t).dt, y=**∫**K(t).P_2_(t).dt. Thus, each dot lies on the *n*-sphere of radius 1 in the space of the *n* principal components of a Principal Component Analysis (PCA). Qualitatively, whereas the filter shape does not change when a mineral oil background is presented (the dots before and after the background presentation are close, Supplementary Figure 5A), the shape changes when a linalool background is presented, Supplementary Figure 5B). We computed the distance between the dots in the PCA space to get a quantitative proxy of the neural dynamics perturbations and compared this proxy between various VPCs. The neural dynamics perturbations appear higher for citronellol, linalyl acetate, Z3HA, and linalool (Supplementary Figure 5C).

In order to assess the quality of the L-N model, we fitted the model to the test dataset and computed the Pearson correlation coefficient between the experimental firing rate and the predicted firing rate (Supplementary Figure 6). We observed a good fit for hexenal, mineral oil and α-pinene, meaning that only three parameters (the phase φ, the threshold, and the gain of the non-linear function) are sufficient to capture the Phe-ORNs response dynamics. For the other VPCs, capturing the Phe-ORNs response dynamics would require more parameters and/or cannot be explained solely by the pheromone stimulus.

### Saliency

We computed the “contrast”, or saliency ^14^ of a Phe-ORN response to a 2-s pheromone sequence by using the following formula:

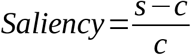

where s is the maximum firing rate during the 2-s pheromone sequence and c is the maximum firing rate during a 2-s window without any pheromone stimulation. For the 2×2s_Phe_ experiment, the maximum firing rate s was examined in the window between 38 s and 40 s (where t=0 s is the onset of the pheromone sequence), and the maximum firing rate was examined in the window between 36 s and 38 s, representing a 2-s pheromone-free interval before the L-2p stimulation.

### Jensen-Shannon divergence

To get a measure of the distance between two Phe-ORNs responses, we computed the Jensen-Shannon divergence (JSD) between two firing rate time courses P and Q using the following formula:

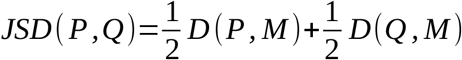

where D is the Kullback-Leibler divergence, and where M=0.5 x (P+Q). The JSD was estimated thanks to the scipy.signal.jensenshannon function in Python. In the 20×2s_Phe_ experiment, we compared the response to the “no background” protocol and the response to the constant or fluctuating protocol, to investigate whether the response to the control resembles the response to a VPC background. In the 2×2s_Phe_ experiment, we compared the response to F-2p and L-2p to investigate whether these two responses are similar.

### Trial-to-trial variability 1: weighted vector strength

To quantify the consistency of the neural responses between repeated trials and whether a VPC can modify the response pattern, we used two complementary approaches: (i) a phase-locking analysis using the weighted vector strength ^82^ and (ii) a spike train dissimilarity analysis ^83^ combined with dimensional reduction by multidimensional scaling for visualization.

Phase locking to the periodic 2-s pheromone stimulus (period T=2 s, 20 repetitions) was quantified using vector strength, which measures the concentration of spikes around a given phase. Each temporal spike is projected onto a circle with a period equal to the duration of the pheromone sequence (2 s) by converting its timing into an angle. For each repetition r, spike times t_r,k_ were converted into phase angles θ_r,k_ according to the following formula:

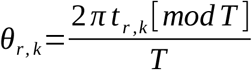

Let Nr be the number of spikes in repetition r. The vector strength V_r_ for each repetition r was computed using the following formula:

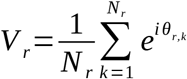

It ranges from 0 (random distribution) to 1 (all spikes aligned in phase). To avoid biases due to adaptation-induced spike count decreases across repetitions, vector strength was computed for each repetition independently, weighted according to the number of spikes, and then averaged across repetitions, so that each trial contributed equally to the final estimate.

To quantify phase-locking consistency across VPC backgrounds, we computed the average vector strength V across the 20 repetitions.

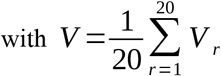

### Trial-to-trial variability 2: spike train temporal dissimilarity

In addition, we assessed the temporal reliability of the responses by calculating the distances between the 20 spike trains, each corresponding to a repetition. For two spike trains *A* and *B*, we defined for each time t the distance to the nearest spike in each train:

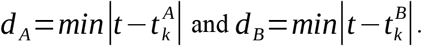

The dissimilarity was defined as:

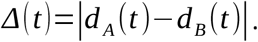

The pairwise spike train distance (RI-SPIKE distance^83^) was computed as the temporal average D(A,B) divided by the inter spike interval, with:

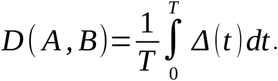

For each background condition (20 repetitions), we computed all pairwise distances, yielding 190 unique distances per background condition.

Distance matrices from the three background conditions were combined into a block matrix, and missing inter-condition distances were replaced by the global mean distance to enable joint embedding in a common coordinate space. The quality of the embedding was assessed using the normalized stress value returned by the Multidimensional Scaling algorithm (MDS, function sklearn.manifold.MDS in Python). For each condition, the 2D MDS coordinates were used to compute the empirical covariance matrix. The 95% confidence ellipse was defined from the eigenvalues of the matrix, scaled by the chi-square quantile. The 95% ellipse does not represent a confidence interval of the mean response but a covariance-based dispersion ellipse, and used as a summary statistic of dispersion rather than as a strict probabilistic boundary. An increase in ellipse area was qualitatively interpreted as a loss of response consistency within a background condition.

### Encoding of pheromone and plant volatile odor pulses

We investigated specifically how Phe-ORNs respond to the presentation of a fluctuating VPC background. Spike trains, pheromone and VPC stimulus values were binned into 1-ms bins for downstream analyses. The response latency computed in the logistic regression analysis was subtracted from the spike timings. We performed generalized linear mixed models to predict the spike train at time t as a function of (1) the pheromone stimulus s_Phe_(t) alone, (2) the VPC stimulus s_VPC_(t) alone, (3) s_Phe_(t) and s_VPC_(t), (4) s_Phe_(t) and s_VPC_(t) and their interaction to unravel potential non-linear effects. The neuron ID was used as a random effect in all models. GLMMs were performed using the package lme4 in R v4.5.0 (R Core Team, 2025). Models were compared based on their respective Akaike Information Criterion (AIC) values and also the relative difference between the effect sizes. The estimated effect of the control stimulus, although statistically significant, corresponds to only a ∼15% increase in spike rate, versus >900% for the pheromone alone — suggesting a negligible functional impact.

### Comparing metrics across different backgrounds

Multiple comparisons of coding metrics across various odor background conditions were performed with R v4.5.0 (packages dplyr, broom, purr, tidyr).

Unless specified, a global ANOVA/Kruskal–Wallis followed by post hoc tests was not used because our hypotheses were restricted to predefined contrasts (comparison between constant and fluctuating backgrounds, comparisons to the solvent control) rather than to all possible pairwise comparisons among VPC conditions, and it would unnecessarily reduce statistical power. For the 20×2s_Phe_ experiment, we compared the metrics between VPCs background and the solvent background (unpaired data), and we compared the metrics in the presence of a fluctuating and a constant background (parametric t-tests or non-parametric Wilcoxon tests). For the 2×2s_Phe_ experiment, we compared the coding performance (or other metrics) to L-2p versus L-20p, and F-2p versus L-2p. We also wanted to compare the coding performance to L-2p across all VPCs. Three sets of planned comparisons (parametric t-tests or non-parametric Wilcoxon tests) were conducted: comparison between fluctuating background versus constant background (paired data), comparison between constant VPC backgrounds and the solvent background (unpaired data), and comparison between VPC backgrounds and the solvent background (unpaired data). False discovery rate correction was applied separately within each family of comparisons. Survival curves were compared using log-rank tests. A linear regression was performed between the firing response to a VPC during the 2-s window preceding the pheromone onset (Figure 1C) and the coding performance thanks to the packages “sklearn.linear_model” and “statsmodel.api” in Python. The package “statsmodel.api” allowed us to compute, for each VPC, the standardized residuals and the Cook’s distance to detect influential points and outliers.

## Acknowledgments

We thank Vincent Jacob for fruitful discussions concerning the linear-nonlinear models. We thank Joanne Louison and Isabelle Touton for *Agrotis ipsilon* rearing. This work was funded by ANR (grant ANR15-CE02-010-01 “Odorscape”).

P.C. benefited from a grant scholarship from the E.N.S. de Lyon.

## Contributions

P.C. and C.M. performed the electrophysiological recordings. P.C. analyzed the experimental data, with inputs from T.B. P.L. designed the stimulus device. P.C. wrote the first draft of the manuscript. M.R. and P.L. supervised the research. All authors critically read the manuscript. The authors declare that they do not have any financial conflicts of interest in relation to the content of the article.

## Supplementary tables

**Supplementary Table 1.** Table of tested VPCs, with their abbreviation used in the text, the CAS number, the dilution deposed on the paper filter, the molar mass and the logP (volatility)

| VPC name | abbreviation | color code | CAS Number | Concentration in source | Concentration in headspace for 1% | Molar mass (g/mol) | logP |
| --- | --- | --- | --- | --- | --- | --- | --- |
| Linalool | Lin | purple | 78-70-6 | 4 | 800μL/20mL | 154.25 | 2.7 - 2.97 |
| (-)-α-pinene | α-p | cyan | 7785-26-4 | 0.75 | 150μL/20mL | 136.23 | 2.8 - 4.83 |
| Eucalyptol | Euc | eucalyptus | 470-82-6 | 2 | 400μL/20mL | 154.25 | 2.5 - 2.74 |
| β-caryophyllene | β-c | light green | 87-44-5 | - | 1 mL/20 mL | 204.35 | 4.4 - 6.30 |
| (Z)-3-hexenyl acetate | Z3HA | orange | 3681-71-8 | 1 | 200μL/20mL | 142.20 | 2.4 |
| (E)-2-hexenyl acetate | E2HA | olive | 2497-18-9 | 1 | 200μL/20mL | 142.20 | 2.4 |
| Linalyl acetate | l-a | pink | 2497-18-9 | 4 | 800μL/20mL | 196.29 | 3.93 |
| Citronellol | Cit | yellow | 106-22-9 | 4 | 800μL/20mL | 156.27 | 3.2 - 3.91 |
| (E)-2-hexen-1-al | Hex | brown | 6728-26-3 | 1 | 200μL/20mL | 98.14 | 1.5 |

**Supplementary Table 2.**
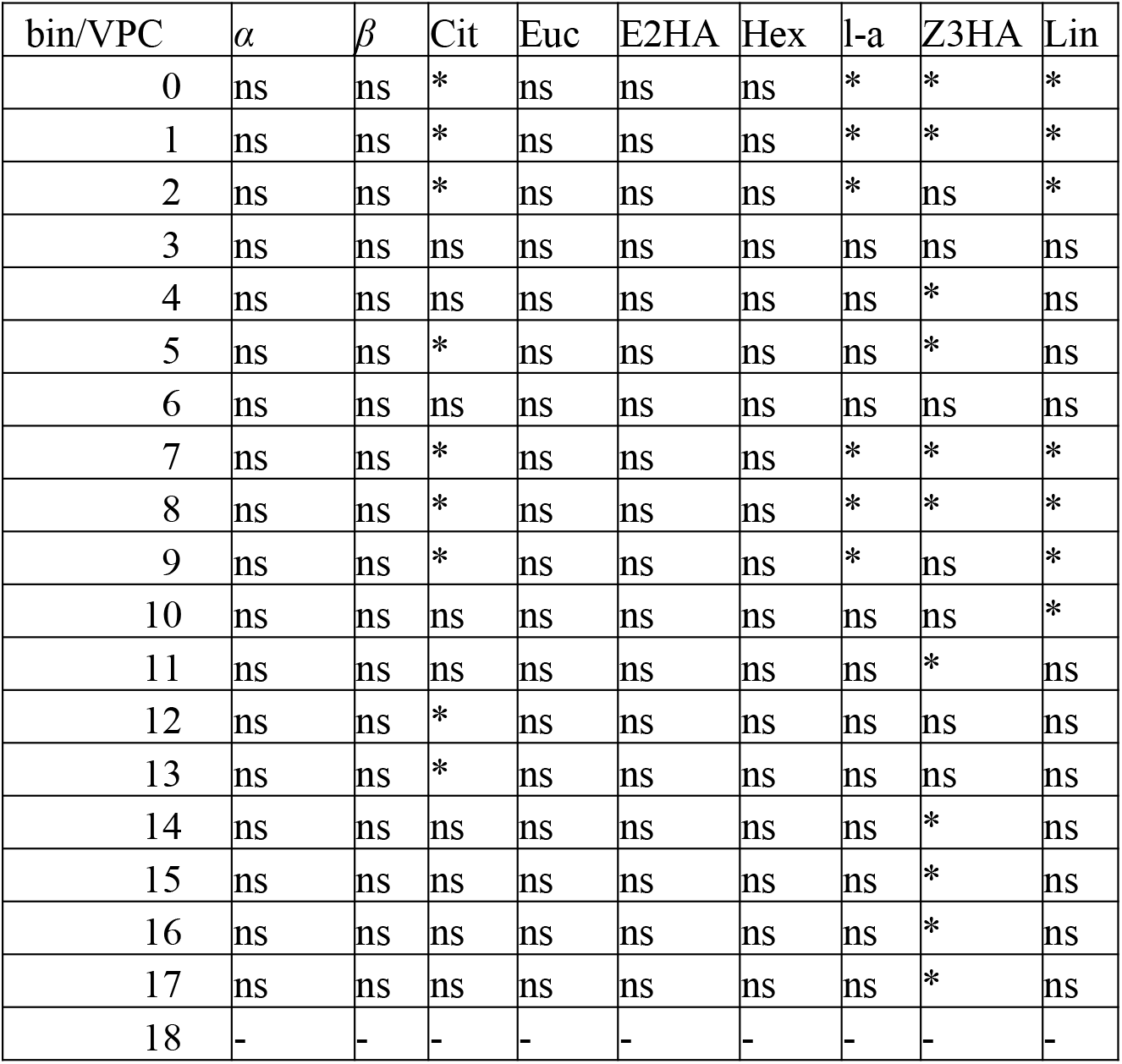
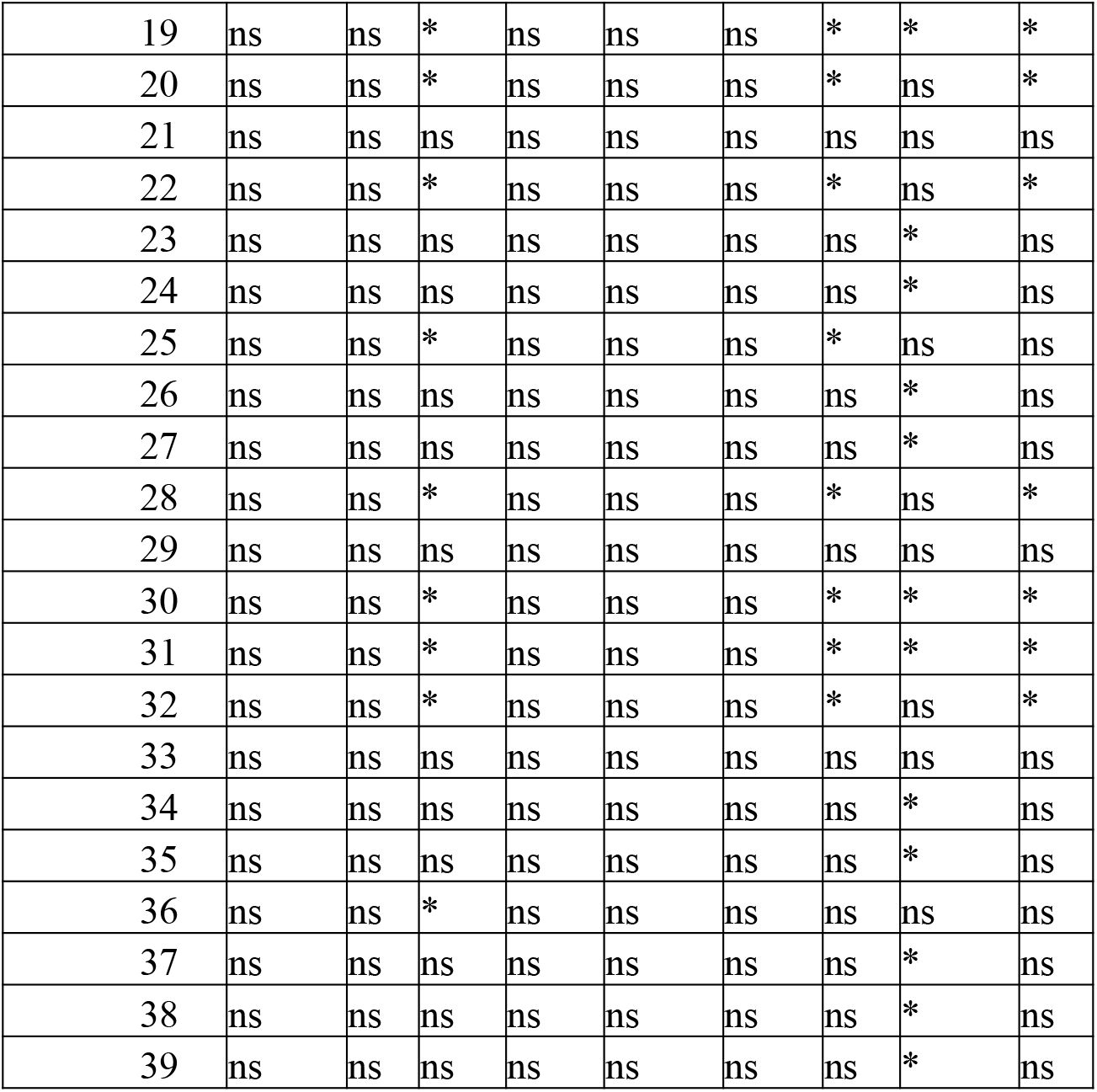
Table of comparisons between the mean error rate for each of the 40 bins present in the repeated 2-s pheromone stimulus.

| bin/VPC | α | β | Cit | Euc | E2HA | Hex | l-a | Z3HA | Lin |
| --- | --- | --- | --- | --- | --- | --- | --- | --- | --- |
| 0 | ns | ns | * | ns | ns | ns | * | * | * |
| 1 | ns | ns | * | ns | ns | ns | * | * | * |
| 2 | ns | ns | * | ns | ns | ns | * | ns | * |
| 3 | ns | ns | ns | ns | ns | ns | ns | ns | ns |
| 4 | ns | ns | ns | ns | ns | ns | ns | * | ns |
| 5 | ns | ns | * | ns | ns | ns | ns | * | ns |
| 6 | ns | ns | ns | ns | ns | ns | ns | ns | ns |
| 7 | ns | ns | * | ns | ns | ns | * | * | * |
| 8 | ns | ns | * | ns | ns | ns | * | * | * |
| 9 | ns | ns | * | ns | ns | ns | * | ns | * |
| 10 | ns | ns | ns | ns | ns | ns | ns | ns | * |
| 11 | ns | ns | ns | ns | ns | ns | ns | * | ns |
| 12 | ns | ns | * | ns | ns | ns | ns | ns | ns |
| 13 | ns | ns | * | ns | ns | ns | ns | ns | ns |
| 14 | ns | ns | ns | ns | ns | ns | ns | * | ns |
| 15 | ns | ns | ns | ns | ns | ns | ns | * | ns |
| 16 | ns | ns | ns | ns | ns | ns | ns | * | ns |
| 17 | ns | ns | ns | ns | ns | ns | ns | * | ns |
| 18 | - | - | - | - | - | - | - | - | - |
| 19 | ns | ns | * | ns | ns | ns | * | * | * |
| 20 | ns | ns | * | ns | ns | ns | * | ns | * |
| 21 | ns | ns | ns | ns | ns | ns | ns | ns | ns |
| 22 | ns | ns | * | ns | ns | ns | * | ns | * |
| 23 | ns | ns | ns | ns | ns | ns | ns | * | ns |
| 24 | ns | ns | ns | ns | ns | ns | ns | * | ns |
| 25 | ns | ns | * | ns | ns | ns | * | ns | ns |
| 26 | ns | ns | ns | ns | ns | ns | ns | * | ns |
| 27 | ns | ns | ns | ns | ns | ns | ns | * | ns |
| 28 | ns | ns | * | ns | ns | ns | * | ns | * |
| 29 | ns | ns | ns | ns | ns | ns | ns | ns | ns |
| 30 | ns | ns | * | ns | ns | ns | * | * | * |
| 31 | ns | ns | * | ns | ns | ns | * | * | * |
| 32 | ns | ns | * | ns | ns | ns | * | ns | * |
| 33 | ns | ns | ns | ns | ns | ns | ns | ns | ns |
| 34 | ns | ns | ns | ns | ns | ns | ns | * | ns |
| 35 | ns | ns | ns | ns | ns | ns | ns | * | ns |
| 36 | ns | ns | * | ns | ns | ns | ns | ns | ns |
| 37 | ns | ns | ns | ns | ns | ns | ns | * | ns |
| 38 | ns | ns | ns | ns | ns | ns | ns | * | ns |
| 39 | ns | ns | ns | ns | ns | ns | ns | * | ns |

**Supplementary Table 3.**
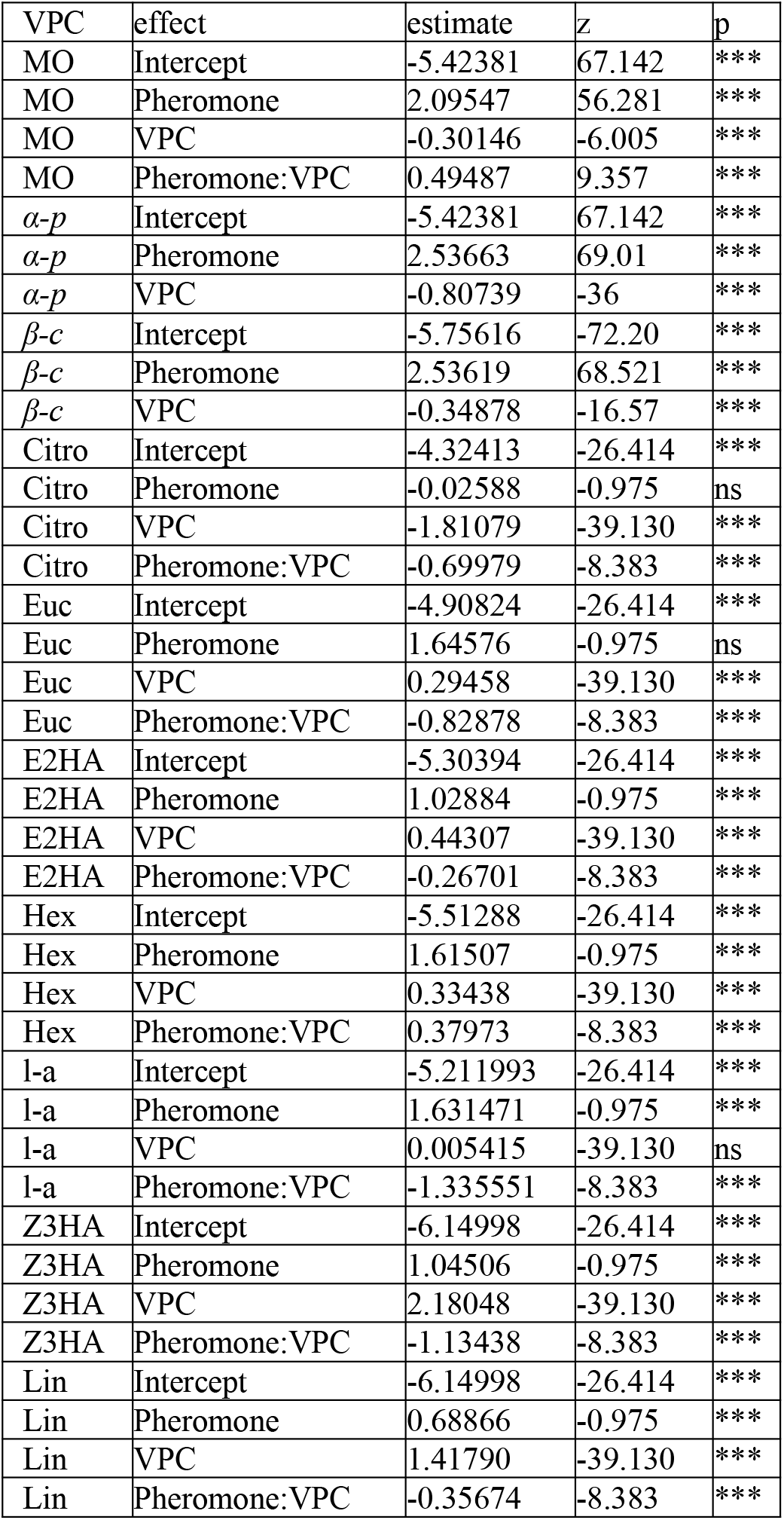
Results of the generalized models used to predict the spike trains as a function of the pheromone and the VPC stimuli.

| VPC | effect | estimate | z | p |
| --- | --- | --- | --- | --- |
| MO | Intercept | -5.42381 | 67.142 | *** |
| MO | Pheromone | 2.09547 | 56.281 | *** |
| MO | VPC | -0.30146 | -6.005 | *** |
| MO | Pheromone:VPC | 0.49487 | 9.357 | *** |
| α-p | Intercept | -5.42381 | 67.142 | *** |
| α-p | Pheromone | 2.53663 | 69.01 | *** |
| α-p | VPC | -0.80739 | -36 | *** |
| β-c | Intercept | -5.75616 | -72.20 | *** |
| β-c | Pheromone | 2.53619 | 68.521 | *** |
| β-c | VPC | -0.34878 | -16.57 | *** |
| Citro | Intercept | -4.32413 | -26.414 | *** |
| Citro | Pheromone | -0.02588 | -0.975 | ns |
| Citro | VPC | -1.81079 | -39.130 | *** |
| Citro | Pheromone:VPC | -0.69979 | -8.383 | *** |
| Euc | Intercept | -4.90824 | -26.414 | *** |
| Euc | Pheromone | 1.64576 | -0.975 | ns |
| Euc | VPC | 0.29458 | -39.130 | *** |
| Euc | Pheromone:VPC | -0.82878 | -8.383 | *** |
| E2HA | Intercept | -5.30394 | -26.414 | *** |
| E2HA | Pheromone | 1.02884 | -0.975 | *** |
| E2HA | VPC | 0.44307 | -39.130 | *** |
| E2HA | Pheromone:VPC | -0.26701 | -8.383 | *** |
| Hex | Intercept | -5.51288 | -26.414 | *** |
| Hex | Pheromone | 1.61507 | -0.975 | *** |
| Hex | VPC | 0.33438 | -39.130 | *** |
| Hex | Pheromone:VPC | 0.37973 | -8.383 | *** |
| l-a | Intercept | -5.211993 | -26.414 | *** |
| l-a | Pheromone | 1.631471 | -0.975 | *** |
| l-a | VPC | 0.005415 | -39.130 | ns |
| l-a | Pheromone:VPC | -1.335551 | -8.383 | *** |
| Z3HA | Intercept | -6.14998 | -26.414 | *** |
| Z3HA | Pheromone | 1.04506 | -0.975 | *** |
| Z3HA | VPC | 2.18048 | -39.130 | *** |
| Z3HA | Pheromone:VPC | -1.13438 | -8.383 | *** |
| Lin | Intercept | -6.14998 | -26.414 | *** |
| Lin | Pheromone | 0.68866 | -0.975 | *** |
| Lin | VPC | 1.41790 | -39.130 | *** |
| Lin | Pheromone:VPC | -0.35674 | -8.383 | *** |

## Supplementary figures

**Supplementary Figure 1.**
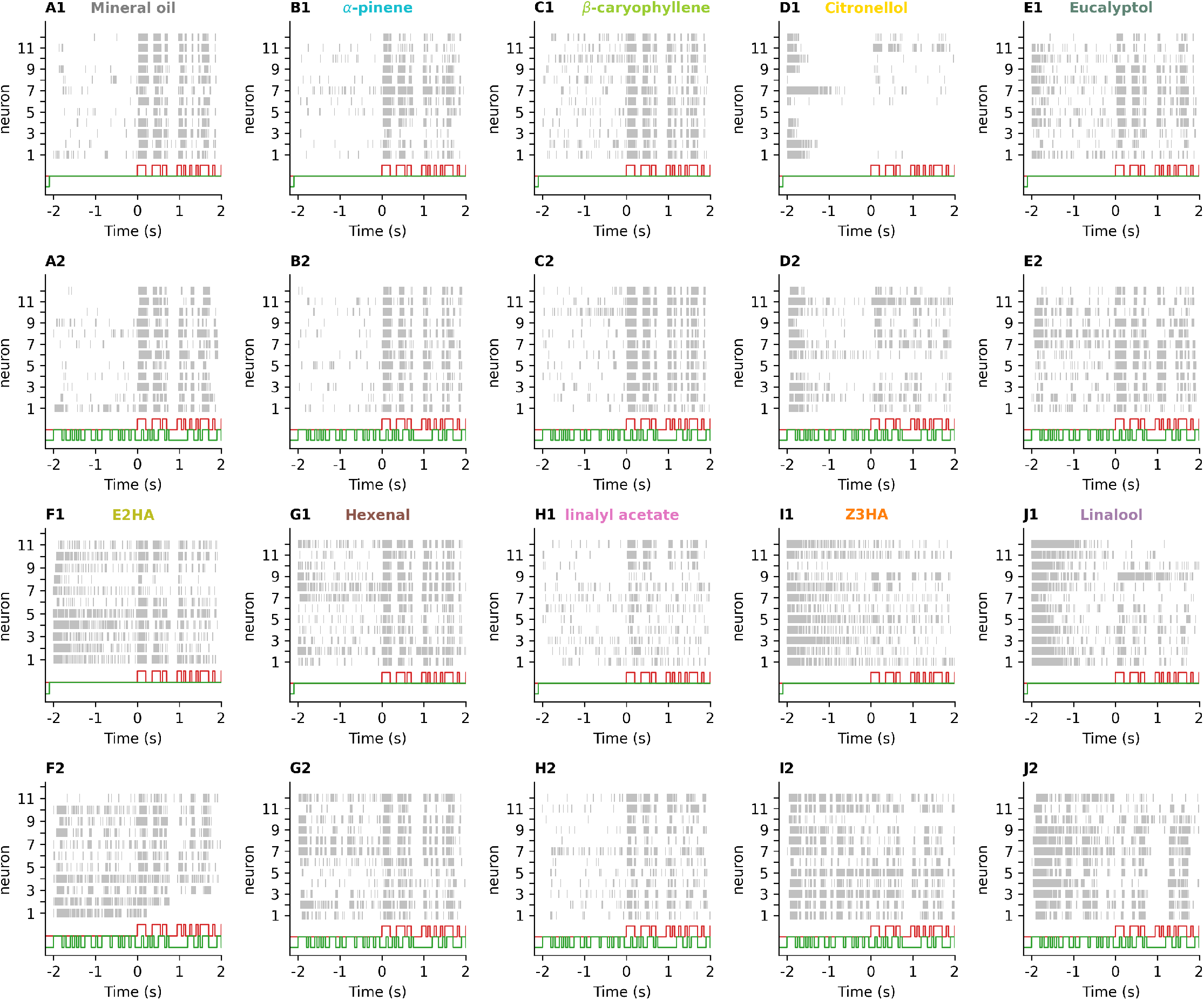
Raster plots of the response of the Phe-ORNs to different VPCs. Raster plots of Phe-ORNs (grey) in the presence of a background of mineral oil (A), α-pinene (B), β-caryophyllene (C), citronellol (D), eucalyptol (E), E2HA (F), hexenal (G), linalyl acetate (H), Z3HA (I) and linalool (J). The pheromone stimulus starts at t=0 s, and the time window shows the response of multiple Phe-ORNs (in grey) two seconds before and after the pheromone stimulus onset. The pheromone stimulus is shown in red, while the VPC stimulus is shown in green. (A1-J1) shows the raster plots of Phe-ORNs to a constant VPC background, while (A2-J2) shows the raster plots of Phe-ORNs to a fluctuating VPC background.

**Supplementary Figure 2.**
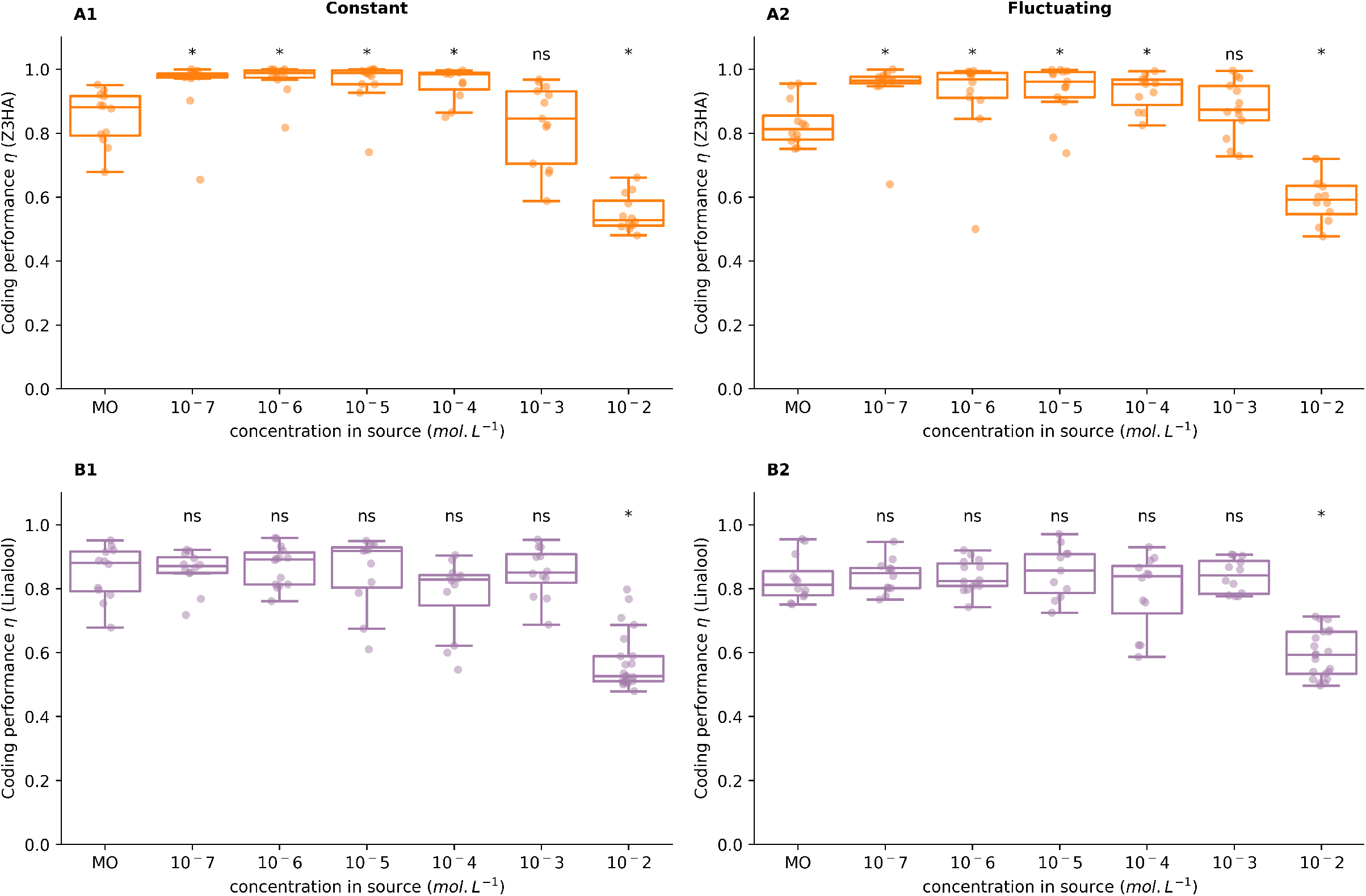
The linalool-induced and Z3HA-induced alterations of coding are dose-dependent. **(A-B)** Boxplots of the temporal coding index η as a function of the concentration of the VPC dilution deposited on the filter paper for (A) Z3HA, in orange, and (B) linalool, in purple, in the presence of a constant (A1, B1) or fluctuating (A2, B2) background. Planned comparisons to mineral oil (MO), ns=non-significant, *: p-value <0.05.

**Supplementary Figure 3.**
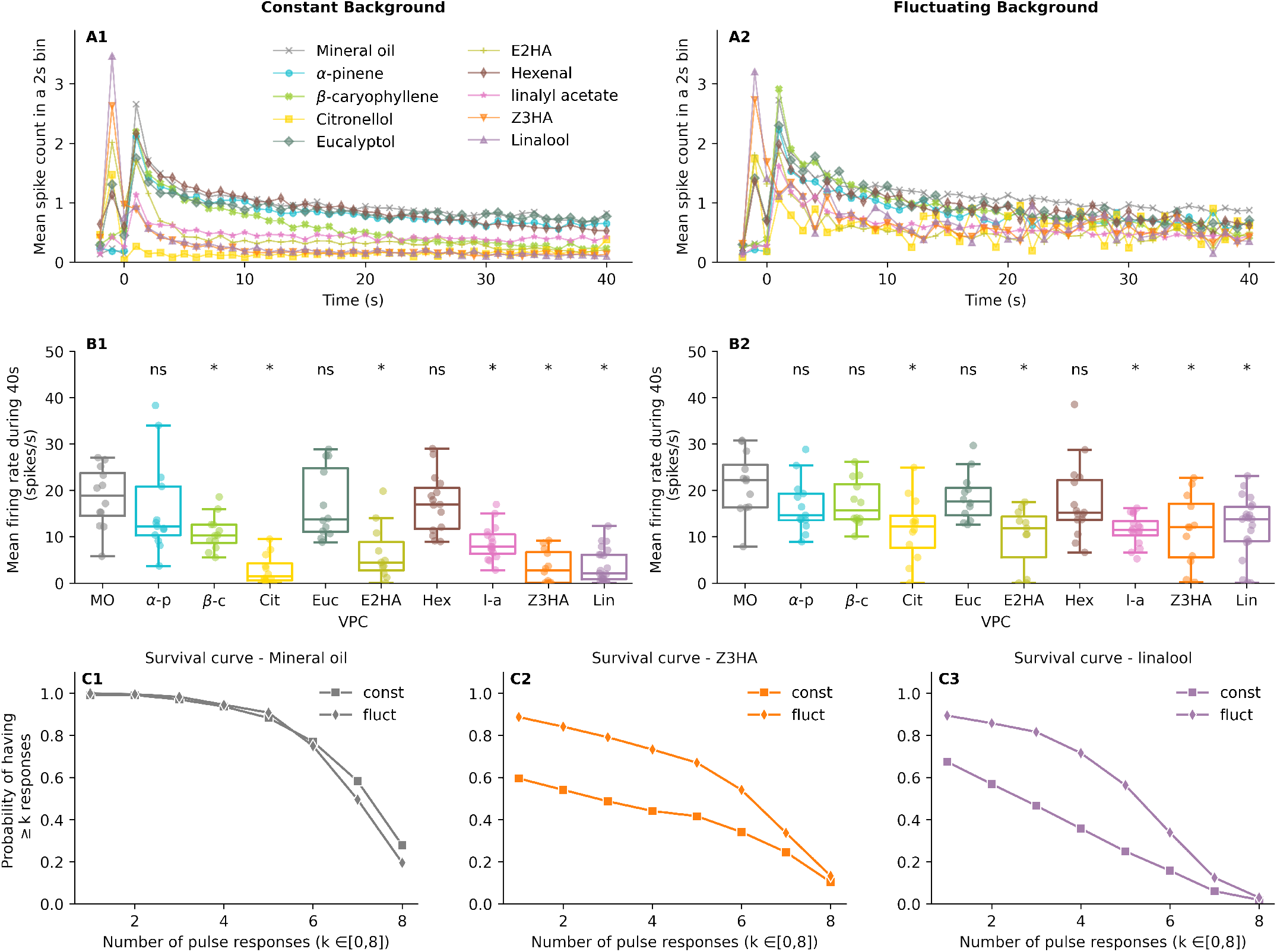
Effects of a VPC background on the mean Phe-ORN response. **(A)**Mean spike count for each of the 2-s bins for a constant (A1) or fluctuating (A2) VPC background. Different VPC backgrounds are shown by different colors. The zero on the x-axis represents the onset of the pheromone stimulus (20 repetitions of the 2-s pheromone sequence, starting 3 s after recording onset, yielding a total of 43 s). Note that some VPCs activate Phe-ORNs (large mean spike count at time t=-1s), whereas a control background (mineral oil, grey) does not activate Phe-ORNs. **(B)**Boxplots of the mean firing rate as a function of the VPC background delivered either as a constant (B1) or fluctuating background (B2). Planned comparisons to mineral oil (MO), ns=non-significant, *: p-value <0.05). **(C)**Survival curves for response persistence (square=constant background, diamond=fluctuating background) for mineral oil (C1), and two VPCs that activate Phe-ORNs: Z3HA (C2) and linalool (C3)

**Supplementary Figure 4.**
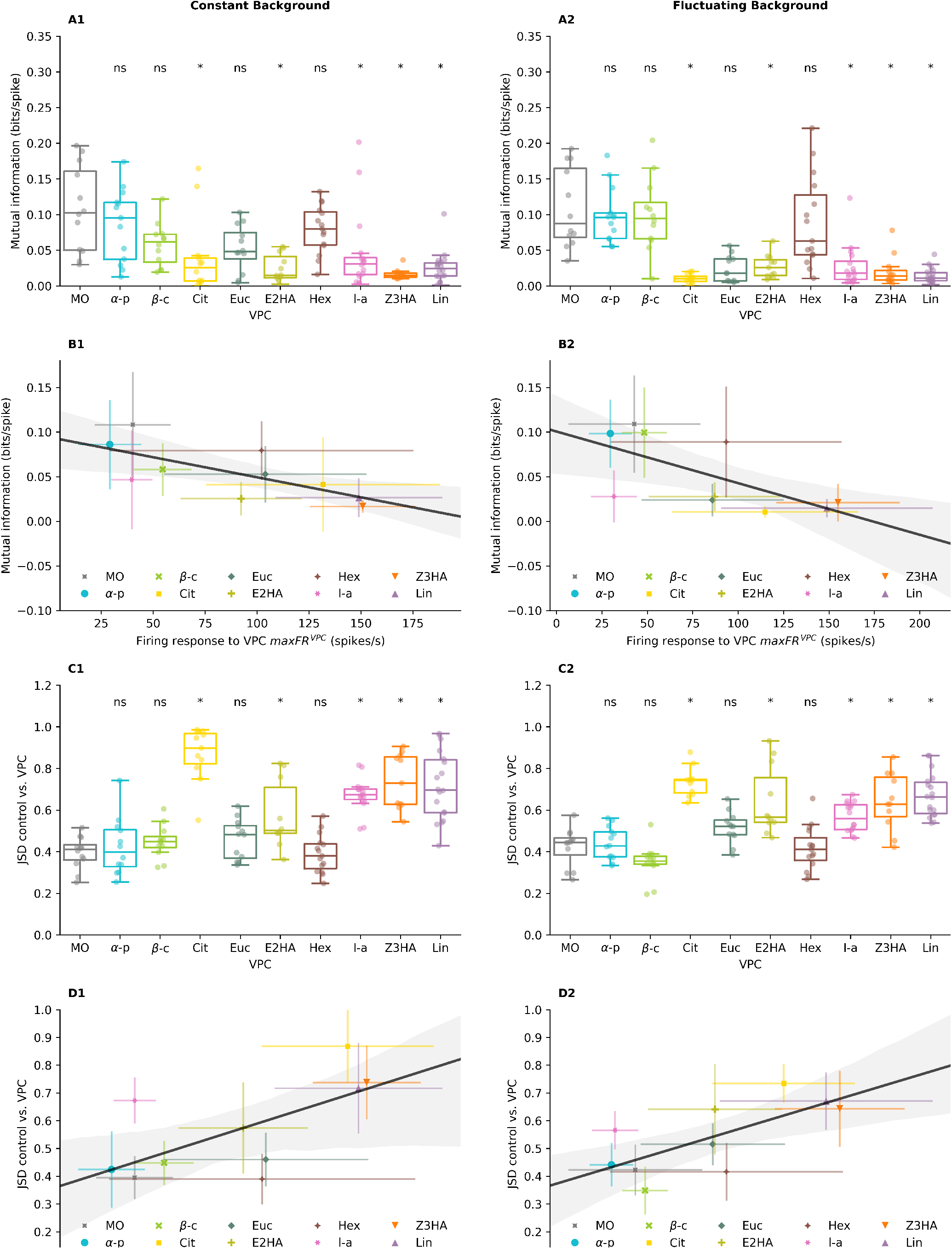
Some VPCs increase the divergence between the Phe-ORNs’ response in clear air and in a VPC background and decrease the mutual information between the spike train and the stimulus. **(A)**Boxplots of the mutual information in the presence of constant (A1) or fluctuating (A2) VPCs background. Planned comparisons to mineral oil (MO), ns=non-significant, *: p-value <0.05. **(B)**Linear regression between the response to a VPC (maximum firing rate during a 2-s window before the pheromone onset) and the mutual information. The fitted regression line is shown in black, and the shaded area indicates the associated confidence interval. **(C)**Boxplots of the JSD (Jensen-Shannon Distance) between the firing rate of the Phe-ORNs in clean air (control) and in the presence of constant (C1) or fluctuating (C2) VPCs background. Planned comparisons to mineral oil (MO), ns=non-significant, *: p-value <0.05. **(D)**Linear regression between the response to a VPC (maximum firing rate during a 2-s window before the pheromone onset) and the JSD. The fitted regression line is shown in black, and the shaded area indicates the associated confidence interval.

**Supplementary Figure 5.**
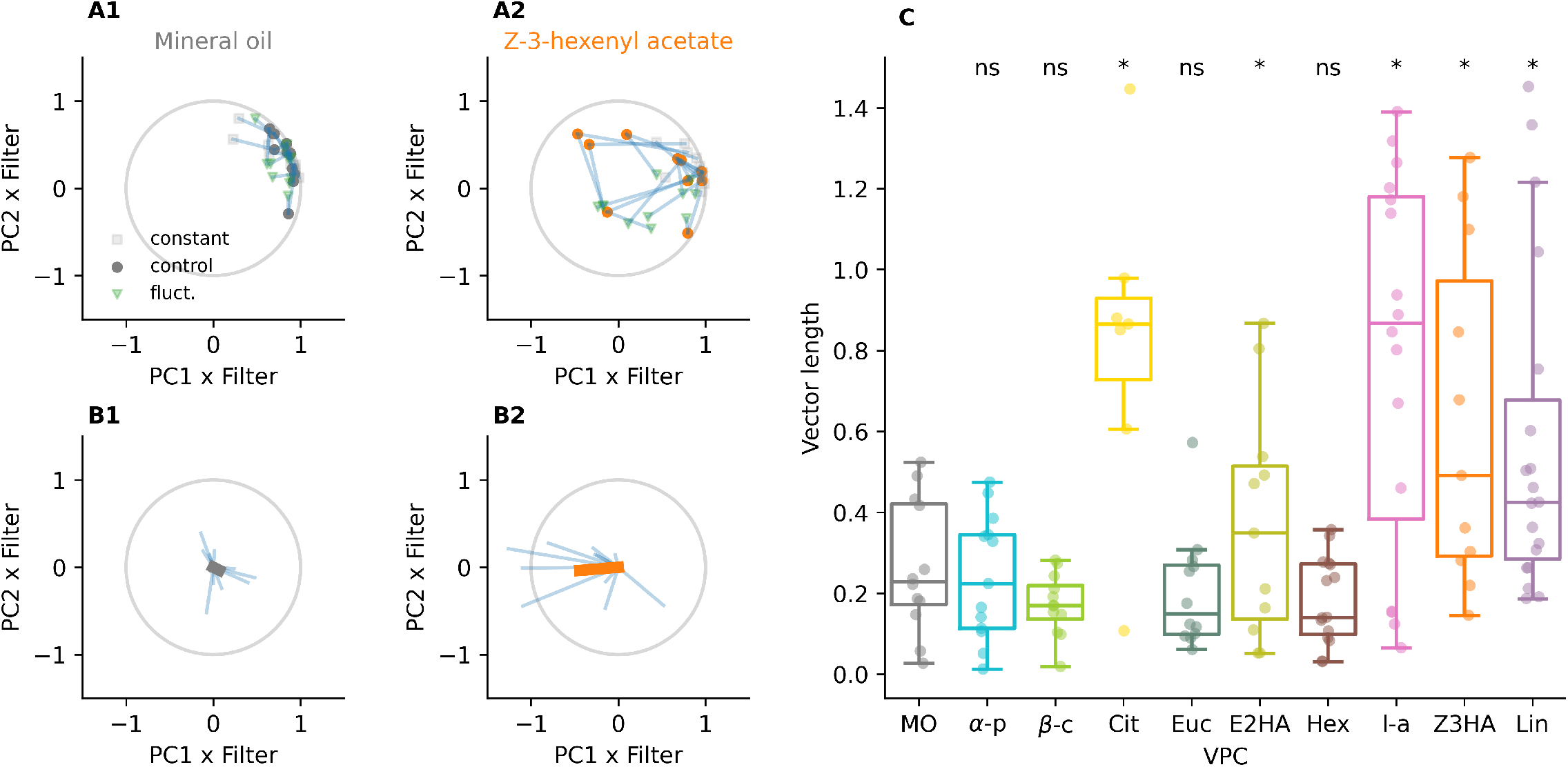
Some VPCs alter Phe-ORN dynamics. **(A-B)** Projection of normalized ORN filters (control, constant background, fluctuating background) on the subspace spanned by the first two components for mineral oil (left column, A1 and B1, grey) or Z3HA (right column, A2 and B2, orange). (A) Each ORN contributes three points, joined by blue lines: the colored circle dot reflects the shape of the filter under the no-background conditions, while the grey square dot and green triangle dot reflect its shape under constant and fluctuating backgrounds, respectively. (B) For each ORN, the difference between the three filters is plotted as a vector from the origin (blue line). The average vector is shown by a colored thick line. **(C)** Boxplots of the vector length between dots in the PCA space (length of the neural “trajectory” between the states ‘no background’, ‘constant background’ and ‘fluctuating background’). Larger vector lengths indicate stronger differences between Phe-ORN responses without or with a VPC background. Planned comparisons to mineral oil (MO), ns=non-significant, *: p-value <0.05.

**Supplementary Figure 6.**
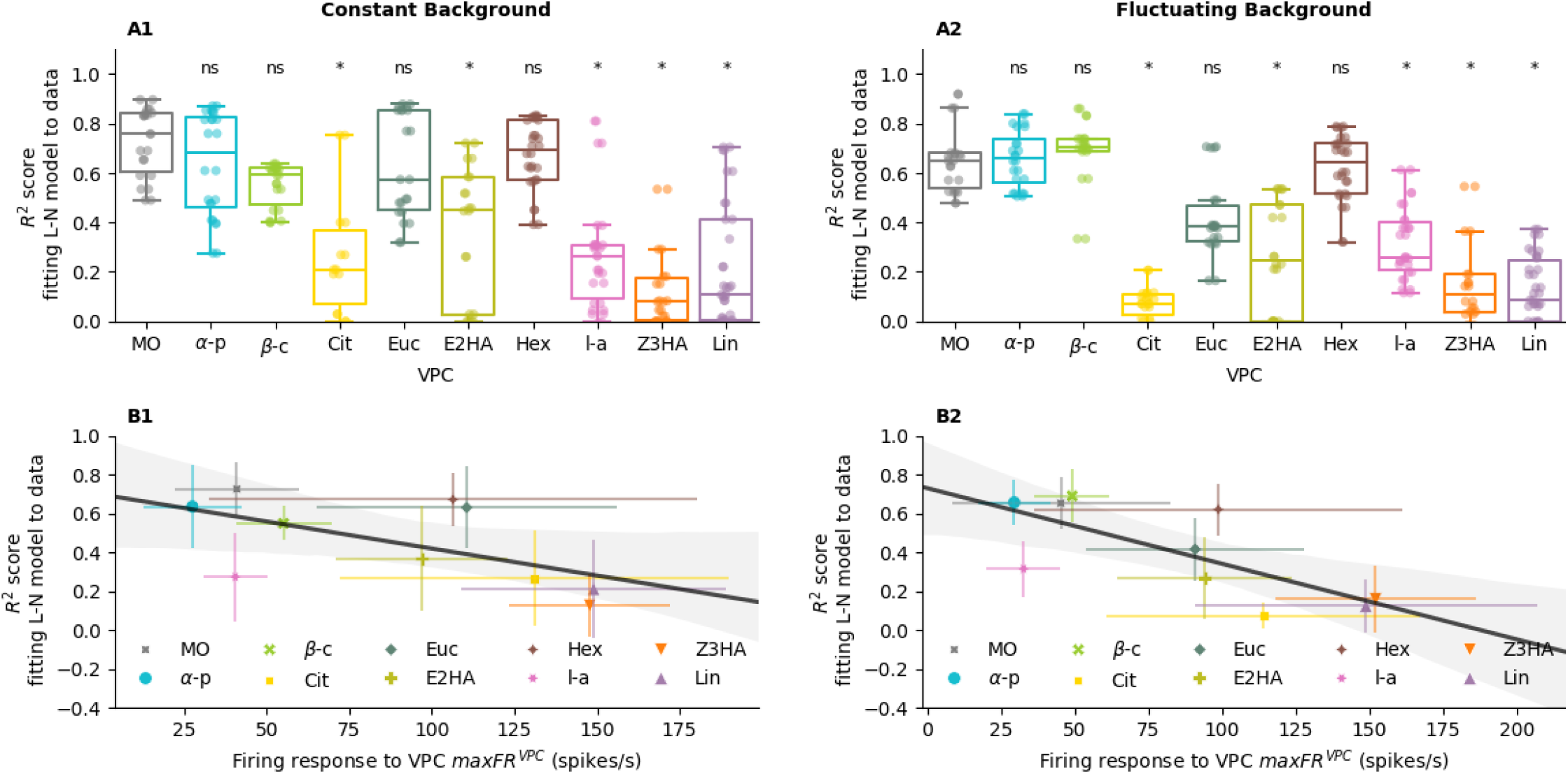
In the presence of some VPCs, more than 3 parameters are needed to capture the Phe-ORN response. **(A)**Boxplots of the correlation coefficient between the L-N model prediction and the experimental firing rate. (Planned comparisons to MO, ns=non-significant, *: p-value <0.05). **(B)**Linear regression between the response to a VPC (maximum firing rate during a 2-s window before the pheromone onset) and the correlation coefficient in (A). The fitted regression line is shown in black, and the shaded area indicates the associated confidence interval.

**Supplementary Figure 7.**
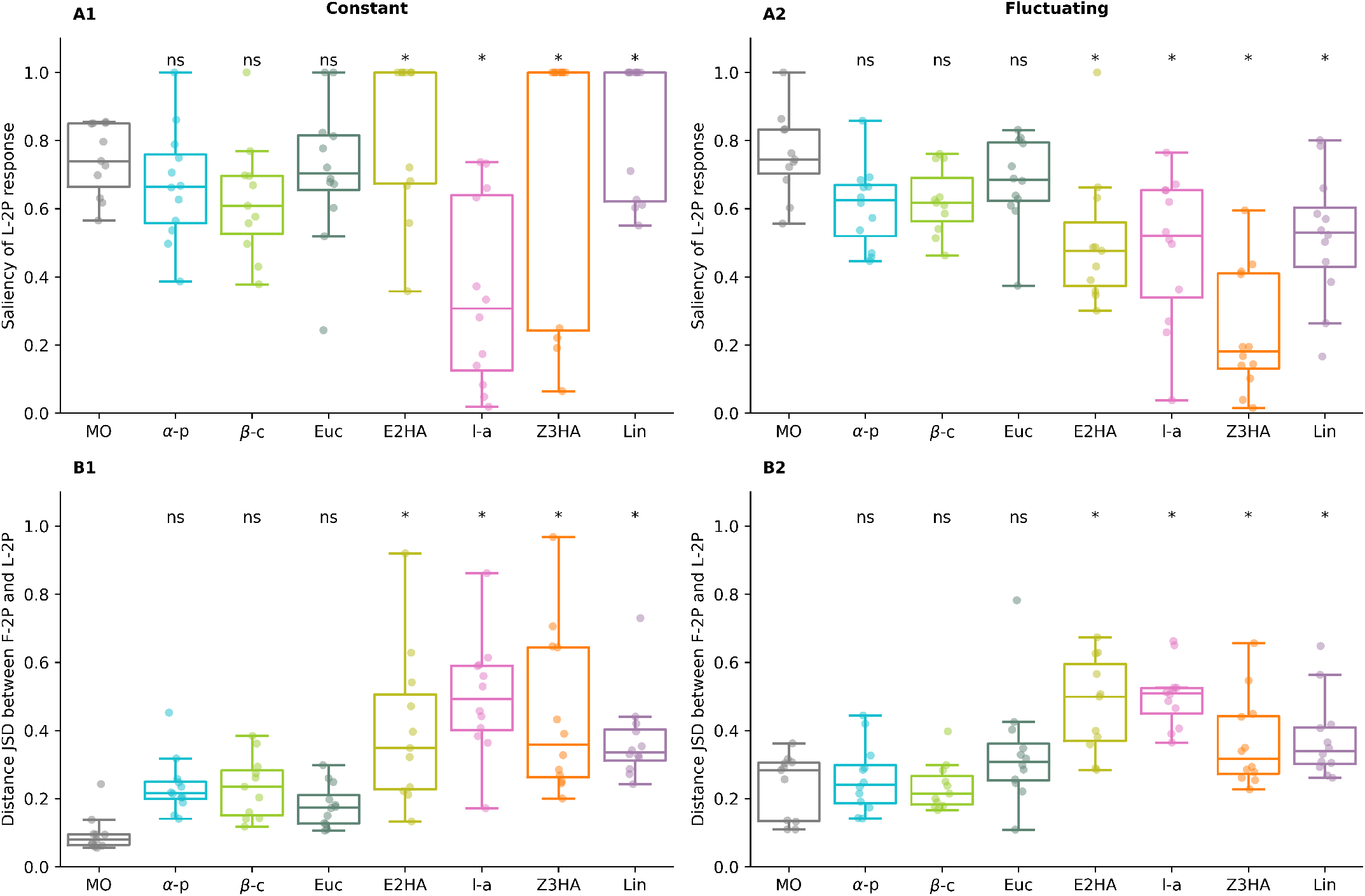
A VPC background reduces the coding performance while increasing the response saliency. **(A)**Saliency of the response to the L-2p as a function of the tested VPCs delivered either as a constant (A1) or intermittent (A2) background. **(B)**Jensen-Shannon Distance between the response to the last 2-s stimulation (L-2p) in the absence versus in the presence of a VPC delivered either as a constant (B1) or fluctuating (B2) background. Planned comparisons to mineral oil (MO), ns=non-significant, *: p-value <0.05.

**Supplementary Figure 8.**
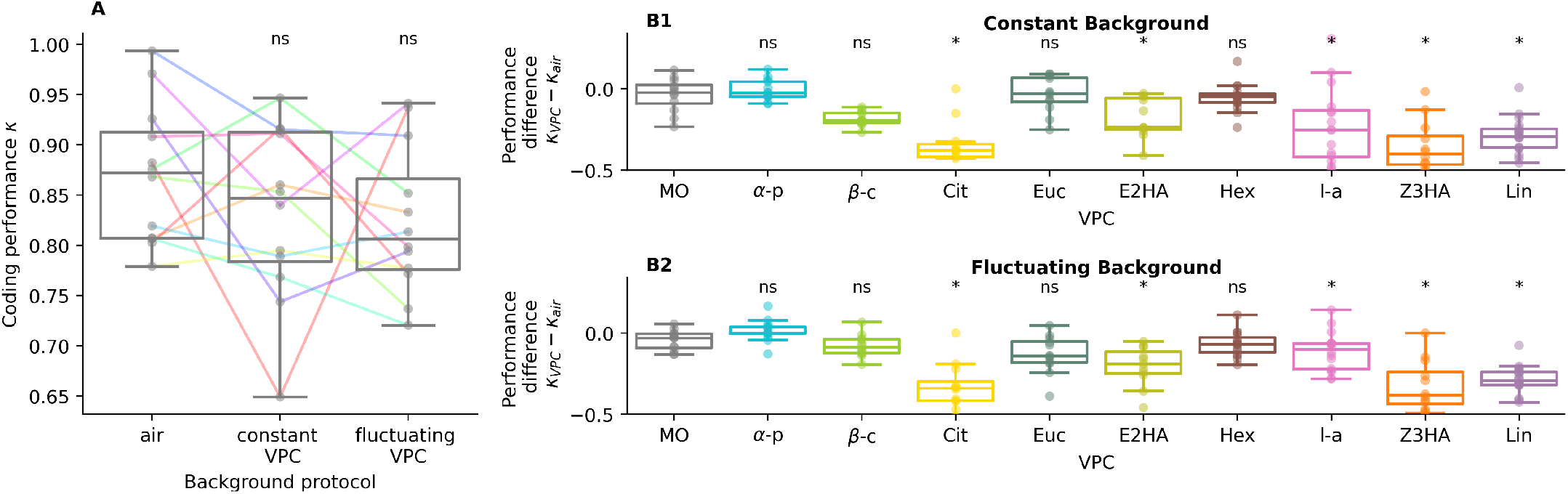
Comparison of the logistic regression -based encoding performance across VPCs, when using a different method. **(A)**We computed the AUC score κ separately for the three stimulation protocols (air, constant VPC background, fluctuating VPC background). Comparison of the performance score κ under a constant mineral oil (solvent) background, a fluctuating mineral oil background or no background. For each neuron, we tested 3 stimulation protocols, thus the data are paired. Colored lines indicate paired data points, and the color code is specific to this subfigure only. (Planned comparisons to ‘air’, ns=non-significant, *: p-value <0.05). **(B)**Difference between the AUC score computed in the presence of a constant (B1) or fluctuating (B2) VPC background (κ_VPC_), minus the AUC score in the absence of a background (κ_air_). (Planned comparisons to MO, ns=non-significant, *: p-value <0.05).

